# ZFP462 targets heterochromatin to transposon-derived enhancers restricting transcription factor binding and expression of lineage-specifying genes

**DOI:** 10.1101/2021.06.28.449463

**Authors:** Ramesh Yelagandula, Karin Stecher, Maria Novatchkova, Luca Michetti, Georg Michlits, Jingkui Wang, Pablo Hofbauer, Carina Pribitzer, Gintautas Vainorius, Luke Isbel, Sasha Mendjan, Dirk Schübeler, Ulrich Elling, Julius Brennecke, Oliver Bell

**Author notes:** Co-first author (Ramesh Yelagandula, Karin Stecher). Correspondence (R.Y), (O.B.).

## Abstract

*ZNF462* haploinsufficiency is linked to Weiss-Kruszka Syndrome, a genetic disorder characterized by a range of neurodevelopmental defects including Autism. Though it is highly conserved in vertebrates and essential for embryonic development the molecular functions of *ZNF462* are unclear. We identified its murine homolog ZFP462 in a screen for epigenetic gene silencing in mouse embryonic stem cells (mESCs). Here, we show ZFP462 safeguards neural lineage specification by targeting the H3K9-specific histone methyltransferase complex G9A/GLP to mediate epigenetic silencing of endodermal genes. ZFP462 binds to thousands of transposable elements (TEs) that harbor ESC- and endoderm-specific transcription factor (TF) binding sites and act as enhancers. Through physical interaction with G9A/GLP, ZFP462 seeds heterochromatin at TE-derived enhancers restricting the binding of core pluripotency TFs OCT4 and SOX2. Loss of ZFP462 in ESCs results in increased chromatin accessibility at target sites and ectopic expression of endodermal genes. Taken together, ZFP462 restricts TF binding and subsequent endoderm-specific gene activation by conferring lineage and locus-specificity to the broadly expressed epigenetic regulator G9A/GLP. Our results suggest that aberrant activation of endodermal genes in the neuronal lineage underlies ZNF462-associated neurodevelopmental pathology.

## Main

Development from a totipotent zygote to a multicellular organism involves numerous cellular specialization steps that are subject to transcriptional and epigenetic regulation^1–3^. Differentiation into divergent cell types is primarily controlled by master regulatory transcription factors (TFs) that bind to regulatory DNA sequences, such as enhancers, and activate lineage-specific genes^4^. However, silencing of lineage non-specific genes is equally important for restricting differentiation to defined pathways. Such silencing involves recruitment of heterochromatin modifiers that restrict DNA accessibility by establishing repressive chromatin modifications including Histone H3 Lysine 9 di- and trimethylation (H3K9me2/3). H3K9me2/3 acts as a binding site for Heterochromatin protein 1 isoforms (HP1)^5^. In turn, chromatin-bound HP1 can oligomerize and recruit additional heterochromatin modifiers, thus promoting chromatin compaction and transcriptional silencing ^6,7^. Epigenetic feedback mechanisms can promote stable inheritance of heterochromatin thereby contributing to the longterm silencing of lineage non-specific genes and maintenance of cell identity ^8–12^.

H3K9me2 modification is catalysed by a heterodimeric complex comprised of G9A (also known as Euchromatic Histone Methyltransferase 2 – EHMT2) and G9-like protein (GLP, also known as Euchromatic Histone Methyltransferase 1 - EHMT1) ^13^. The HMTase complex is essential for embryonic development ^14^. G9A/GLP-dependent heterochromatin modifications at promoters and enhancers have been proposed to block transcription factor binding, thereby preventing unscheduled activation of lineage non-specific gene expression ^15–18^. Indeed, G9A/GLP-dependent heterochromatin silences the pluripotency-linked *Oct3/4* (also known as *Pou5f1*) gene during embryonic stem cell (ESC) differentiation and is critical for preventing its expression in somatic tissues ^19,20^. Inactivation of G9A facilitates *Oct3/4* gene induction during iPSC reprogramming of somatic cells, supporting the importance of this H3K9-specific HMTase in limiting developmental potential ^21^. G9A/GLP has been proposed to function as a master regulator of neurodevelopment by restricting neuron-specific gene expression to the nervous system ^22^. G9A/GLP interaction with RE1-silencing transcription factor (REST also known as NRSF) targets and represses neuronal genes in nonneuronal cells ^23–25^. However, G9A/GLP also has important and distinct functions in silencing both non-neuronal and early neuron progenitor genes in mature neurons ^26,27^. Notably, *de-novo* mutations in genes encoding either GLP itself or proteins that cooperate in facultative heterochromatin formation cause Kleefstra syndrome phenotypic spectrum, a neurodevelopmental disorder that is characterized by intellectual disability, autism-like phenotypes, childhood hypotonia, and craniofacial dysmorphology ^28^.

Together, this suggests that G9A/GLP-dependent heterochromatin plays an important role in establishment and maintenance of lineage-specific expression patterns during development. While the biochemical functions and developmental importance are well-established, the molecular mechanisms underlying precise spatiotemporal silencing of lineage non-specific genes by G9A/GLP remain largely unclear ^13,29,30^. Since G9A/GLP are broadly expressed, and lack sequence-specificity, heterochromatin mediated by the H3K9me2-specific HMTase complex ultimately relies on celltype specific interactions with TFs which are currently poorly defined.

Using a CRISPR genetic screen for modifiers of heterochromatin-mediated silencing of *Oct3/4*, we identified Zinc finger protein 462 (ZFP462), a vertebrate-specific, putative transcription factor of unknown function. Notably, its human ortholog, *ZNF462*, has recently been identified as a novel, high - confidence risk gene for a neurodevelopmental disorder (OMIM#: 618619) ^31–33^. Here, we demonstrate that ZFP462 functions as transcription factor required for silencing of inappropriate endodermal gene expression in mouse embryonic stem cells (mESCs) and neural progenitor cells (NPCs). ZFP462 controls epigenetic silencing of lineage non-specific genes by recruiting of G9A/GLP to repress transposon (TE)-derived enhancers. ZFP462 specifically directs G9A/GLP to form heterochromatin at these TE-derived enhancers thereby limiting DNA accessibility for aberrant binding of pluripotency TFs. Hence, our work demonstrates that ZFP462 precisely targets formation of G9A/GLP-dependent heterochromatin to prevent unscheduled activation of endodermal genes and to maintain cell fate identity. Our findings provide insight into the developmental regulation of G9A/GLP-dependent gene silencing. In addition, our data suggest that neural phenotypes associated with *ZNF462* haploinsufficiency are linked to defects in cell fate specification at early steps of embryonic development.

## Results

### *Zfp462* encodes a novel regulator of H3K9me2/3-mediated gene silencing

To dissect the molecular mechanisms underlying heterochromatin-mediated silencing of a well-defined developmental target gene, we focused on the *Oct3/4* gene where developmental regulation by heterochromatin is well-documented ^19,21,34^. The Oct4 transcription factor is highly expressed in mESCs and essential for pluripotency and self-renewal ^35^. Upon differentiation, the *Oct3/4* gene promoter is rapidly inactivated and acquires heterochromatin modifications including H3K9me2. Through subsequent HP1 binding and DNA methylation, expression of *Oct3/4* is irreversibly silenced ^20,21,34^. Experimental tethering of the chromo shadow domain of HP1 (HP1) to a genetically modified *Oct3/4* allele in mouse embryonic stem cells (mESCs) is sufficient to target H3K9-specific histone methyltransferases and recapitulates heterochromatin-mediated gene silencing^12^. We modified this original chromatin in vivo assay (CiA) at *Oct3/4* to perform a CRISPR-based screen in mESCs. Seven Tet operator DNA binding sites (TetO) and a Puromycin-BFP reporter gene were placed downstream of the *Oct4-GFP* reporter allele enabling efficient, reversible HP1 tethering via the Tet-OFF system (Fig. 1a). After establishing a stable cell line expressing hCas9, we generated reporter cell clones expressing HP1 fused to a FLAG-Tet repressor domain (TetR-FLAG-HP1) (Extended Data Fig. 1a). TetR binds to TetO sites with high affinity but this interaction is fully reversible upon Doxycycline (Dox) addition (Fig. 1b). TetR-FLAG-HP1 expression, but not TetR-FLAG alone, resulted in transcriptional silencing of the *Oct4-GFP* reporter gene (Fig. 1c and Extended Data Fig. 1b). Reporter gene silencing coincided with loss of active histone modifications (H3K4me3) and formation of heterochromatin marked by H3K9me2 and H3K9me3 modifications (Fig. 1d). In addition, the synthetic heterochromatin domain was bound by endogenous HP1, in agreement with previous results (Extended Data Fig. 1c)^12^. After Dox-dependent release of HP1 tethering, transcriptional silencing was robustly maintained through genome replication for more than eight cell divisions (four days) (Fig. 1c). Chromatin Immunoprecipitation (ChIP) followed by qPCR analysis confirmed that heterochromatic histone modifications persist in the absence of the initial stimulus (Fig. 1d and Extended Data Fig. 1c). Notably, H3K9me2 remained highly enriched after eight days of Dox treatment suggesting that ectopic heterochromatin recapitulates epigenetic silencing of the endogenous *Oct3/4* gene in somatic cells by the G9A/GLP complex^19^.

**Fig. 1:**
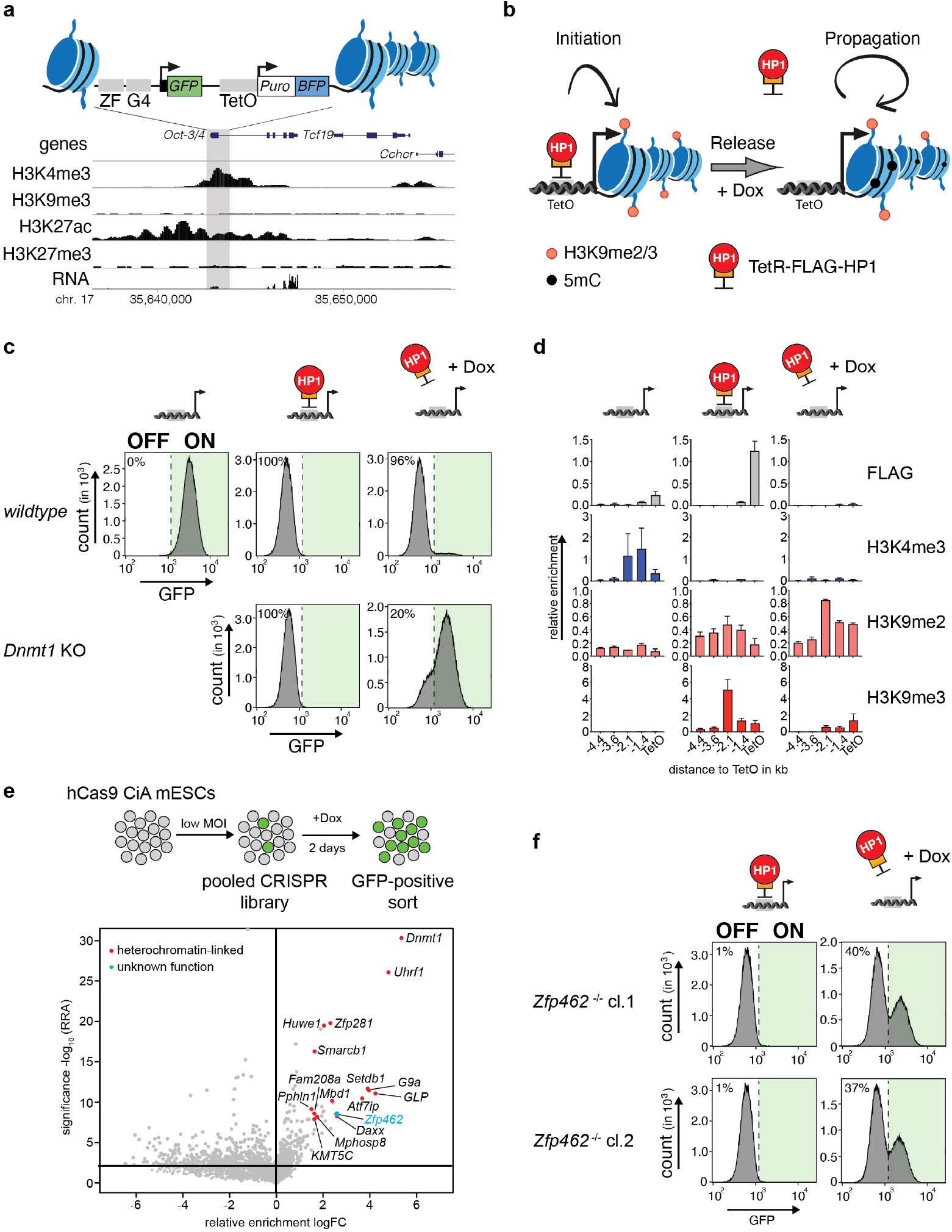
CRISPR screen identifies heterochromatin regulators required for heritable *Oct3/4* gene silencing. **a)** Design of the CiA *Oct4* dual reporter locus in mESCs. One allele of *Oct3/4* was modified in mESCs by inserting seven Tet Operator sites (TetO) flanked by a GFP and a BFP reporter gene on either side. GFP expression is under control of the *Oct3/4* promoter whereas a PGK promoter drives BFP expression. The genomic screen shot (below) shows histone modifications and RNA expression at the *Oct3/4* locus. **b)** Scheme of the experimental design. TetR facilitates reversible HP1 tethering to TetO binding sites to establish heterochromatin and silence both GFP and BFP reporters. Doxycycline (Dox) addition releases TetR binding to distinguish heritable maintenance of chromatin modifications and gene silencing in the absence of the sequencespecific stimulus. **c)** Flow cytometry histograms of wildtype and *Dnmt1* KO CiA *Oct4* dual reporter mESCs show GFP expression before TetR-FLAG-HP1 tethering, in the presence of TetR-FLAG-HP1 and after four days of Dox-dependent release of TetR-FLAG-HP1. Percentages indicate fraction of GFP-negative cells. **d)** ChIP-qPCR shows relative enrichment of TetR-HP1 (FLAG) and histone modifications surrounding TetO before TetR-FLAG-HP1 tethering, in the presence of TetR-FLAG-HP1 and after four days of Dox-dependent release of TetR-FLAG-HP1. Data are mean ± SD (error bars) of at least two independent experiments. **e)** Scheme of CRISPR screen design. MOI refers to multiplicity of infection. Volcano plot shows enrichment (log fold change GFP-pos. sorted vs unsorted cells) and corresponding significance (-log10 MAGeCK significance score) of genes in CRISPR screen (n = mean of three independent experiments). **f)** Flow cytometry histograms show GFP expression of two independent *Zfp462 ^-/-^* CiA *Oct4* dual reporter cell lines in the presence of TetR-FLAG-HP1 and after four days of Dox-dependent release of TetR-FLAG-HP1. Percentages indicate fraction of GFP-positive cells.

We next tested the sensitivity of heterochromatin-mediated epigenetic silencing to genetic perturbation. DNA methyltransferase inhibition using 5-azacytidine impairs mitotic inheritance but not initiation of HP1-dependent reporter gene silencing^12^. To probe the genetic dependence of epigenetic memory on DNA methylation, we infected the TetR-HP1 reporter cell clones with lentiviral vectors expressing sgRNA specific for DNA methyltransferase 1 (*Dnmt1*). Loss of global DNA methylation in *Dnmt1* mutant CiA mESC clone was confirmed by LC-MS (Extended Data Fig. 1d). As expected, heterochromatin-dependent reporter gene silencing was established but failed to be maintained upon Dox treatment of *Dnmt1* mutant CiA mESCs (Fig. 1c). Thus, our modified CiA reporter mESCs recapitulate stable inheritance of heterochromatin and are sensitive to genetic perturbation offering a unique entry point to interrogate the mechanism of cooperation between H3K9me2/3 and DNA methylation.

To identify genes impacting heterochromatin-dependent silencing of *Oct3/4*, we performed a pooled CRISPR screen with unique molecular identifiers (UMIs), which allows analysis of mutant phenotypes at a single-cell level ^36^. The UMI CRISPR library contained approx. 27,000 sgRNAs targeting all annotated mouse nuclear protein-coding genes with four sgRNAs per gene. Each sgRNA was paired with thousands of barcodes representing UMIs improving signal-to-noise ratio and hit calling. hCas9-expressing CiA reporter cells were transduced with the pooled library, selected for five days, and treated with Dox for an additional two days before isolating GFP-positive cells by FACS (Fig. 1e). The unsorted population served as background control. Relative enrichment of sgRNAs was determined by sequencing of UMIs in both populations followed by statistical analysis using MAGeCK^37^. 130 genes were significantly enriched in the GFP-positive cell population (P-value <0.001) (Fig. 1e and Supplementary Table 1). Among the top hits were *Dnmt1, Uhrf1, Setdb1, G9a, GLP, Atf7ip, Daxx, Atrx, KMT5C* and genes encoding members of the HUSH complex, all of which have been previously linked to DNA methylation or H3K9me2/3-dependent gene silencing^13,38,39^. The uncharacterized *Zfp462* gene had not been associated with heterochromatin regulation but scored very highly in the screen. We validated loss of heritable *Oct4-GFP* silencing in *Zfp462* mutant CiA mESCs by independent CRISPR-Cas9 targeting using two distinct sgRNAs against *Zfp462* followed by flow cytometry analysis (Fig. 1f and Extended Data Fig. 1e).

Together, our CRISPR screen revealed several regulators of H3K9me2/3 and DNA methylation supporting previous reports of crosstalk between these two chromatin modification pathways^12,38^. In addition, we uncovered *Zfp462* whose role in heterochromatin regulation we set out to further explore.

### ZFP462 elicits transcriptional silencing through interaction with G9A/GLP and HP1

To investigate the molecular function of ZFP462, we engineered mESCs by inserting an Avi-GFP-FLAG tag into the endogenous *Zfp462* gene (*Avi-Zfp462*) (Fig. 2a). ZFP462 interacting proteins were identified using affinity purification under stringent conditions (300mM NaCl) in mESCs, followed by liquid chromatography mass spectrometry (LC-MS). Consistent with a function of ZFP462 in maintenance of heterochromatin-mediated gene silencing, we uncovered protein interactions with several well-known co-repressors (Fig. 2b and Supplementary Table 2). Most significantly, ZFP462 was strongly associated with HP1, G9A, GLP and WIZ, a known G9A/GLP interacting protein^40^. ZFP462 robust binding to the G9A/GLP complex in mESCs was corroborated using proteomic analysis of affinity purified Avi-fused endogenous GLP (Avi-GLP). This assay revealed enrichment of ZFP462 and HP1 in addition to the canonical GLP interaction partners, G9A and WIZ (Fig. 2c and Supplementary Table 3)^40^. These results argue that ZFP462 functions in transcriptional silencing as it interacts with co-repressor proteins involved in heterochromatin regulation. Since G9A/GLP are ubiquitously expressed and lack tissue- and sequence-specificity, we hypothesized that ZFP462 serves as a tissue- and sequence-specific transcription factor that targets the histone methyltransferase complex to specific genes in defined lineages.

**Fig. 2:**
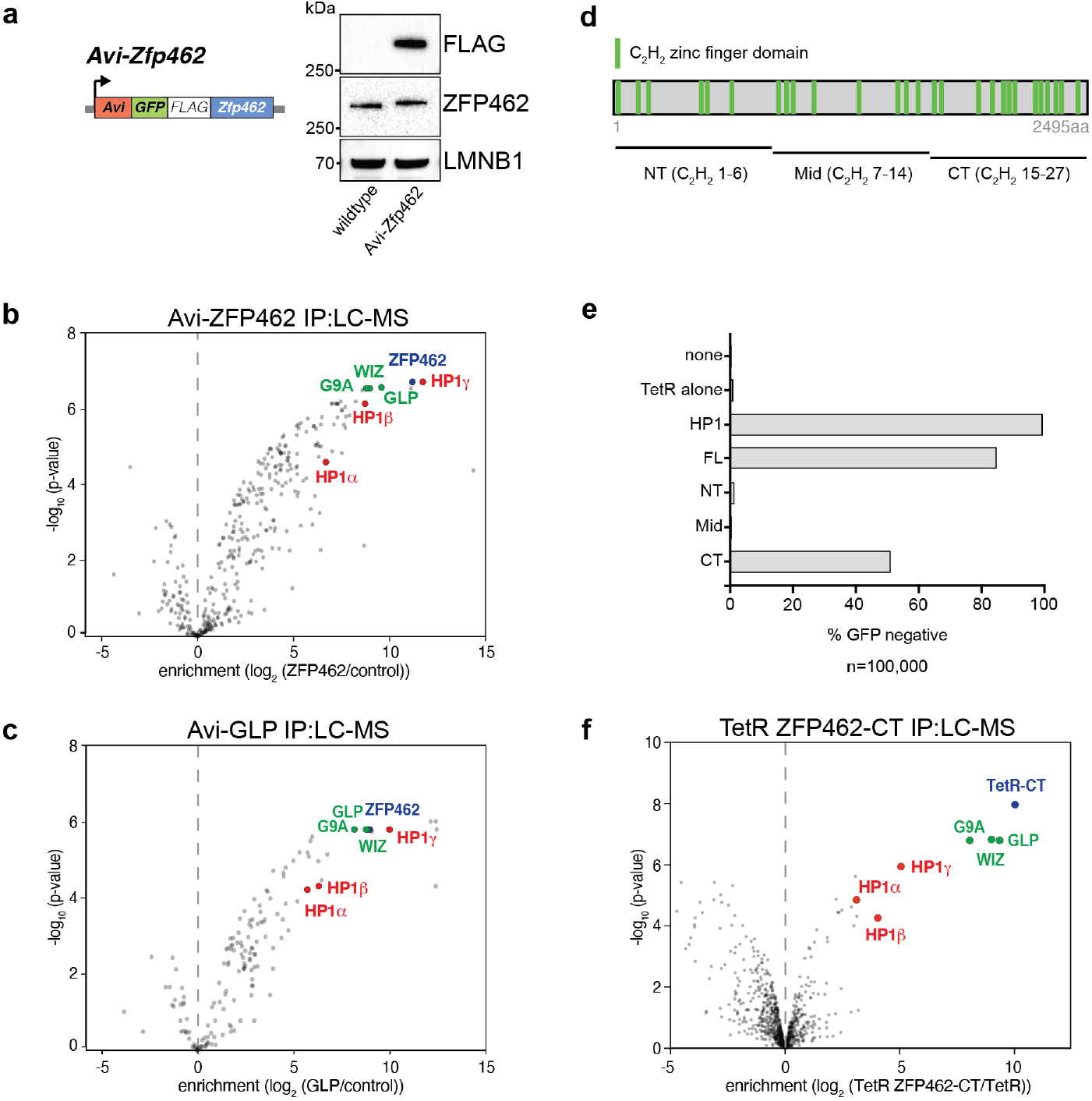
ZFP462 elicits silencing function through interaction with G9A/GLP and HP1. **a)** Design of Avi-*Zfp462* mESCs and western blot validation. Mouse ESCs expressing Biotin ligase (BirA) were used to modify the endogenous *Zfp462* gene by inserting the Avi-GFP-3XFLAG tag downstream of the translation start codon. Western blot with FLAG and ZFP462 antibodies confirms ZFP462 tagging. **b)** and **c)** LC-MS analysis of Avi-tagged ZFP462 and Avi-tagged GLP mESCs. Volcano plots show enrichment and corresponding significance of co-purified proteins. (n = three replicates). **d)** Scheme of ZFP462 protein depicts locations of 27 C2H2 zinc finger domains (green bars). Fragments used to generate TetR fusions for tethering in CiA *Oct4* dual reporter assay are indicated below. **e)** Bar plot shows percentage of GFP-negative CiA *Oct4* mESCs measured by flow cytometry in response to ectopic TetR fusion protein expression (y-axis). **f)** Volcano plot of LC-MS analysis compares enrichment and corresponding significance of co-purified proteins between TetR-FLAG-ZFP462-CT and TetR-FLAG (n = three replicates).

To test this model, we used CiA mESCs to determine if ectopic ZFP462 tethering was sufficient to induce GFP reporter gene silencing. We further examined which protein domains are required for co-repressor interactions. ZFP462 is a highly conserved, vertebrate-specific C2H2-type zinc finger protein (Extended Data Fig. 2a). It contains 27 C2H2 zinc fingers distributed throughout the protein (Fig. 2d). C2H2-type zinc fingers are typically associated with DNA binding activity but may also contribute to protein-protein interactions^40,41^. We transduced CiA mESCs with TetR-FLAG fusions carrying full-length ZF462 (FL), or N-terminal (NT), mid (Mid) and C-terminal (CT) fragments (Fig. 2d and Extended Data Fig. 2b). TetR-mediated targeting of fulllength ZFP462 led to loss of GFP expression in more than 80% of CiA reporter cells supporting a role in recruitment of transcriptional corepressors (Fig. 2e and Extended Data Fig. 2c). Similarly, tethering of the CT fragment resulted in substantial GFP silencing suggesting that C2H2-type zinc fingers 15 through 27 function in corepressor binding. In contrast, GFP silencing was not observed upon recruitment of ZFP462 NT- and Mid fragments. Finally, to determine the nature of repression, we immunoprecipitated the TetR-FLAG CT fusion and performed LC-MS analysis, which revealed strong enrichment of G9A, GLP, WIZ and HP1 (Fig. 2f, Supplementary Table 4). In conclusion, ZFP462 associates with G9A/GLP and HP1 via its C-terminal domain and is sufficient to initiate transcriptional gene silencing.

### ZFP462 represses primitive endoderm differentiation

To determine the biological role of ZFP462, we generated homozygous *Zfp462* mutant mESCs using CRISPR-Cas9 (*Zfp462 ^-/-^*). In addition, we engineered premature stop codons into one allele of *Zfp462* using CRISPR-Cas9 assisted homology-dependent repair (*Zfp462 ^+/Y1195*^* and *Zfp462 ^+/R1257*^*). In particular, *Zfp462 ^+/R1257*^* mutant mESCs mimic human ZNF462 haploinsufficiency associated with Weiss-Kruzska Syndrome (*ZNF462 ^+/R1263*^*)^31^. We isolated two independent *Zfp462* mutant clones for each genotype and confirmed reduction and loss of ZFP462 protein by western blot (Fig. 3a) and heterozygous point mutation by sanger sequencing (Extended Data Fig. 3a). Compared to wildtype mESCs, heterozygous and homozygous *Zfp462* mutant mESCs displayed morphological changes with dispersed, refractile cells spreading out of characteristically densely packed mESC colonies (Fig. 3b and Extended Data Fig. 3b). Further, *Zfp462* mutant mESC colonies showed reduced expression of the pluripotency marker alkaline phosphatase, suggesting a function for ZFP462 in the maintenance of mESC self-renewal. Transcriptome profiling by RNA-seq revealed robust differential expression of ~1800 genes in homozygous *Zfp462* mutant mESCs and ~1400 genes in heterozygous *Zfp462* mutant mESCs, respectively (cutoff: LFC +/-1, padj. 0.01) (Fig. 3c and Extended Data Fig. 3c-d). Gene Ontology terms of both up- and downregulated genes were related to developmental processes, consistent with the spontaneous mESC differentiation we observed upon *Zfp462* deletion (Fig. 3e, Extended Data Fig. 3e). Notably, several key developmental transcription factors were among the upregulated genes in both heterozygous and homozygous *Zfp462* mutant mESCs, including *Bmp4, Gata6, Gata4*, and *Sox17*, required for cell fate specification towards meso- and endodermal lineages (Fig. 3c-d). *Gata6* and *Sox17* expression is restricted to primitive endoderm in early embryos, and overexpression of these transcription factors (TFs) in mESCs is known to promote primitive/extraembryonic endoderm-like differentiation^42–44^. The resulting cells have a similar morphology and gene expression profile to heterozygous and homozygous mutant *Zfp462* mESCs^42^. These results argue that *Zfp462* is required for maintenance of mESC pluripotency and *Zfp462* haploinsufficiency causes the upregulation of key developmental genes including TFs that promote meso-endodermal lineage-specification.

**Fig. 3:**
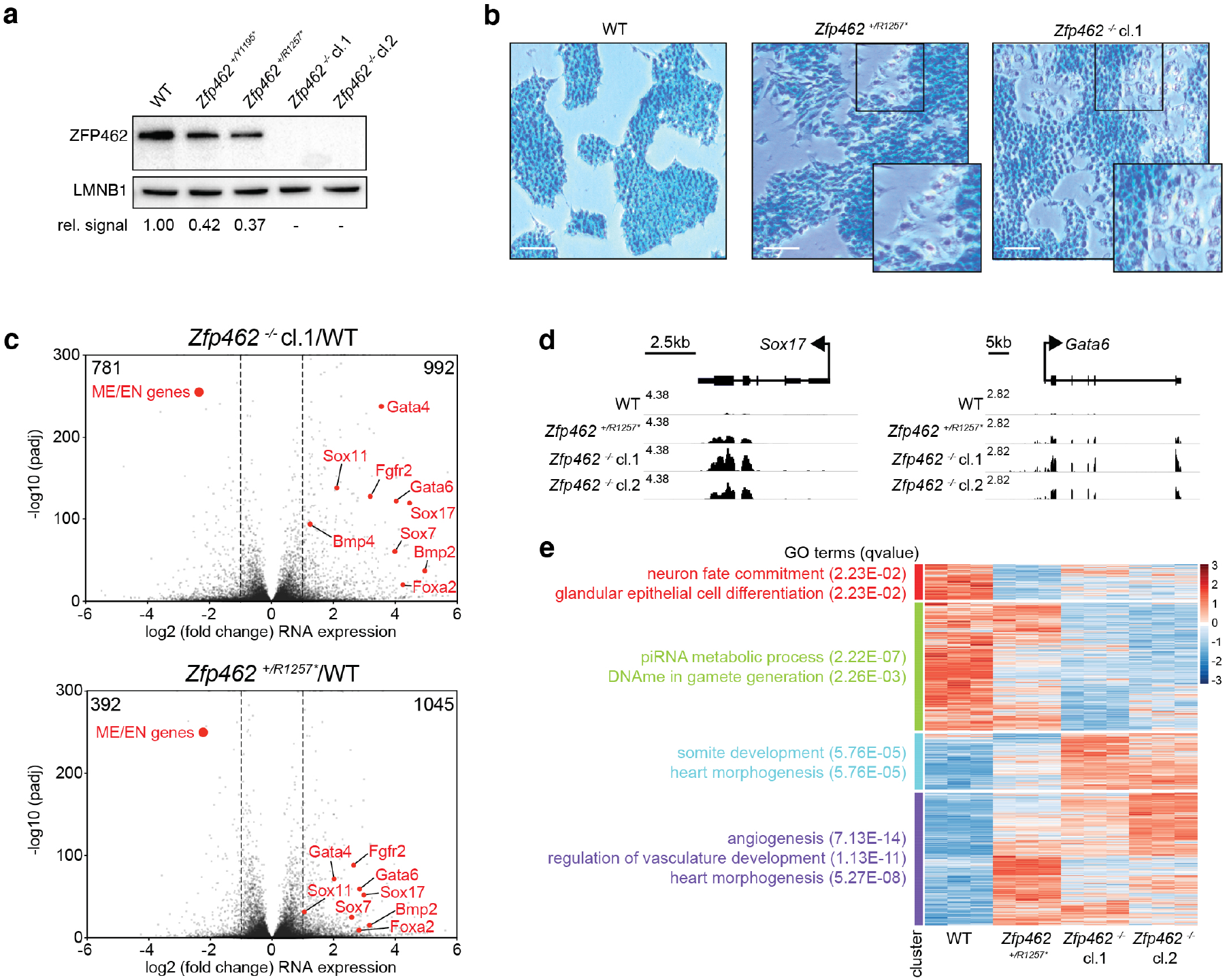
Depletion of *Zfp462* leads to aberrant expression of lineage specifying genes. **a)** Western blot shows ZFP462 protein expression in wildtype (WT), two heterozygous and two homozygous *Zfp462* mutant mESC lines. LMNB1 serves as loading control and reference for relative ZFP462 quantification (numbers below). **b)** Alkaline phosphatase staining of WT, heterozygous and homozygous *Zfp462* mutant mESCs. Enlarged region is marked as square in the image. Scale bar = 100 μm. **c)** Volcano plots show gene expression changes in homozygous (top) and heterozygous (bottom) *Zfp462* mutant mESCs compared to WT mESCs (n = three replicates). Indicated are the numbers of significantly up- or down regulated genes. (padj. = 0.05; LFC = 0.5). **d)** Genomic screen shots of *Sox17* and *Gata6* show mRNA expression levels in WT and *Zfp462* mutant mESCs. All RNA-seq profiles are normalized for library size. **e)** Heatmap show cluster analysis of differentially expressed genes (padj. = 0.05; LFC 1) in WT and *Zfp462* mutant mESCs. Top gene ontology (GO) terms and corresponding significance are indicated for each cluster (left).

### ZFP462 restricts expression of endodermal-lineage genes during neural differentiation

Given the link between *Zfp462/ZNF462* haploinsufficiency and defects in mouse and human neurodevelopment^31,32,45^, we investigated how heterozygous and homozygous *Zfp462* deletions influenced neural differentiation relative to wildtype mESCs. To reduce variability prior to differentiation, we tested if enforcing the naïve ground state using inhibitors against mitogen-activated protein kinase (MAPK) and glycogen synthase kinase-3 (GSK3) (termed 2i) could restore self-renewal of *Zfp462* mutant mESCs. Culturing *Zfp462* mutant mESCs under 2i conditions (2i/S/L) suppressed spontaneous differentiation yielding homogenous, compact colonies similar to wildtype mESCs (Fig. 4a and Extended Data Fig. 4a-b). Consistent with reduced ZFP462 dependence, naïve ground state mESCs grown in 2i/S/L medium displayed lower ZFP462 expression levels (Extended Data Fig. 5a-b). We therefore induced neuronal differentiation starting from naïve ground state mESCs using an established protocol^46^. Sequential withdrawal of 2i and LIF initiated mESC differentiation into embryoid bodies (EBs) containing progenitors of ecto-, meso- and endodermal lineages. Subsequent addition of retinoic acid stimulated ectoderm expansion followed by enrichment of neuronal progenitor cells (NPCs). ZFP462 expression levels increased during neural differentiation with highest levels in NPCs, in agreement with previous reports (Extended Data Fig. 5a-b)^45^. Consistent with its role in neurodevelopment, heterozygous and homozygous *Zfp462* mutants gave rise to EBs and NPCs with reduced size compared to wildtype mESCs (Fig. 4a, Extended Data Fig. 4b). Nevertheless, RT-qPCR analysis revealed that *Nanog* and *Oct4* were downregulated and *Pax6* and *Ngn2* were upregulated, indicating successful exit from pluripotency and induction of neural differentiation in wildtype and mutant cells (Fig. 4b). Expression of endodermal markers *Gata6* and *FoxA2* is strongly induced in *Zfp462* KO mESCs upon 2i withdrawal and remained high even in NPCs (Fig. 4b). Although heterozygous *Zfp462* mutant cells showed similar marker gene expression levels to wildtype cells at the onset of differentiation, they failed to downregulate endoderm-specific genes in NPCs similar to KO cells.

**Fig. 4:**
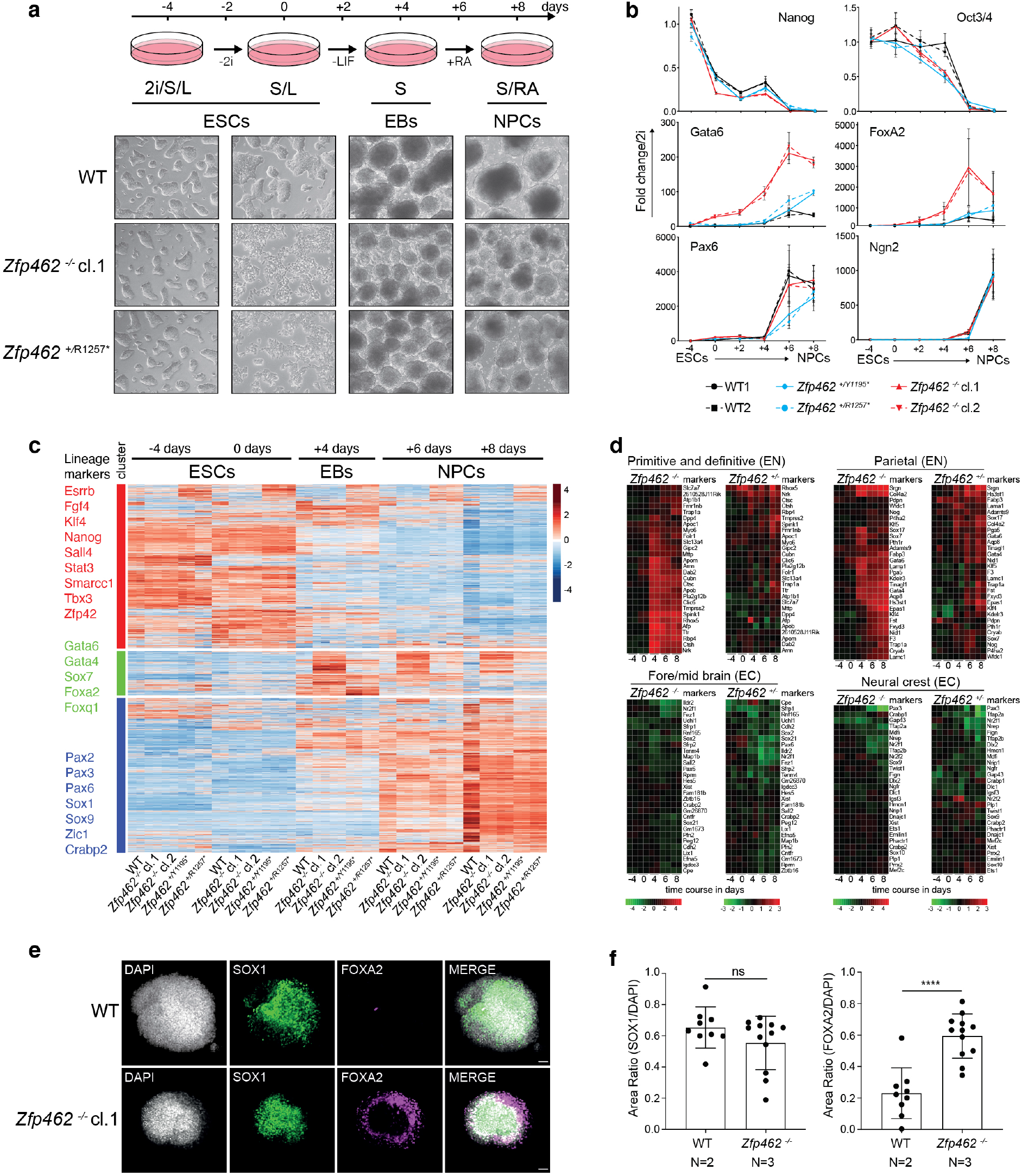
*Zfp462* mutant cells show abnormal cell fate specification during neuronal differentiation. **a)** Scheme shows design of neural differentiation experiment from mESCs to neural progenitor cells (NPCs) (top). Stepwise withdrawal of 2i inhibitors (2i) and Leukaemia Inhibitory Factor (LIF) leads to formation of cellular aggregates called embryoid bodies (EBs). Subsequent treatment with retinoic acid (RA) induces enrichment of NPCs. Representative bright field images show WT and *Zfp462* mutant cells at corresponding stages of neural differentiation. **b)** Line plots shows RT-qPCR analysis of lineage marker expression during neural differentiation (n=two replicates). Expression levels are shown relative to mESCs (2i/S/L). **c)** Heatmap shows cluster analysis of differentially expressed genes (padj. = 0.05; LFC 1) in WT and *Zfp462* mutant cells during neural differentiation (n = two replicates). Selected lineage marker genes are indicated for each cluster (left). **d)** Heatmaps show differential expression of selected marker genes specific for endodermal and neural lineages in heterozygous and homozygous *Zfp462* mutant cells during neural differentiation. **e)** Immunohistochemistry analysis shows SOX1 and FOXA2 expression in WT and homozygous *Zfp462* mutant at day 8 of neural differentiation. Cell aggregates are counterstained with DAPI. Scale bar = 50μm. **f)** Bar plots show quantification of SOX1 and FOXA2 immunofluorescence in WT and *Zfp462 ^-/-^* cell aggregates (n = two independent experiments for WT and three independent experiments for *Zfp462 ^-/-^*).

To gain more comprehensive insight into aberrant gene regulation, we profiled global gene expression changes throughout the differentiation time course (Fig. 4c). Cluster analysis of differentially regulated genes yielded three separate groups based on distinct expression kinetics. Genes in cluster 1 comprising mESC-specific markers, were highly expressed in mESCs under 2i/S/L and S/L conditions and were downregulated in EBs and NPCs. Cluster 2 contained ~1300 genes including meso- and endodermal lineage markers. These genes were induced in EBs and gradually decreased in NPCs derived from wildtype mESCs. In contrast, in heterozygous and homozygous *Zfp462* mutant cells cluster 2 genes were highly induced in EBs and remained overexpressed in NPCs (Fig. 4c-d and Extended Data Fig. 4c). Addition of RA induced upregulation of cluster 3 genes including key neural TFs. Compared to wildtype cells, upregulation of neural genes was lower in *Zfp462* mutant NPCs suggesting a delay in neural differentiation and/or maturation (Fig. 4c-d and Extended Data Fig. 4c). Although deregulation of meso- and endodermal genes was less severe, neural gene expression was similarly impaired in heterozygous and homozygous mutant cells. This is consistent with significant neurodevelopmental defects seen in patients with *ZNF462* haploinsufficiency^31,32^. Finally, we used immunohistochemistry in NPCs which revealed that the deregulation of neuronal and endodermal lineage genes extended to the protein level (Fig. 4e-f). Most wildtype NPCs were SOX1-positive whereas FOXA2 expression was barely detected, in agreement with efficient neural differentiation^47^. In contrast, enrichment and distribution of FOXA2-positive cells was strongly increased in *Zfp462* KO NPCs. Together, these results indicate that *Zfp462* deletion results in misspecification towards the endodermal lineage under conditions that normally induce neural cell identity. Hence, ZFP462 is required to silence expression of lineage non-specific genes and promotes specification into neural cell lineages. These findings are consistent with developmental delay and defects in neural development in heterozygous *Zfp462* KO mice^45^.

### ZFP462 targets heterochromatin through sequence-specific recruitment of G9A/GLP

To investigate how ZFP462 contributes to lineage specification, we used Chromatin Immunoprecipitation followed by sequencing (ChIP-seq) to profile binding of Avi-tagged ZFP462 genome-wide. This analysis revealed 16,259 significantly enriched sites (Fig. 5a, Extended Data Fig. 6a). Most ZFP462 peaks (86%) were found in intergenic and intronic regions, distal to transcription start sites. Since the related family of Krüppel-associated box (KRAB) domain-containing zinc finger proteins (KZFP) target heterochromatin modifiers to silence transposable elements (TE)^48^, we analyzed the overlap of ZFP462 binding with repeat DNA families. This uncovered a strong enrichment of ZFP462 binding at mouse-specific long terminal repeat (LTR) families of endogenous retroviruses (ERVs) (Fig. 5b). Specifically, ZFP462 frequently occupied RLTR9a, RLTR9d, RLTR9d2, RLTR9a2 and RLTR9e families, suggesting a role in repression of these TE subfamilies by G9A/GLP recruitment (Fig. 5c).

**Fig. 5:**
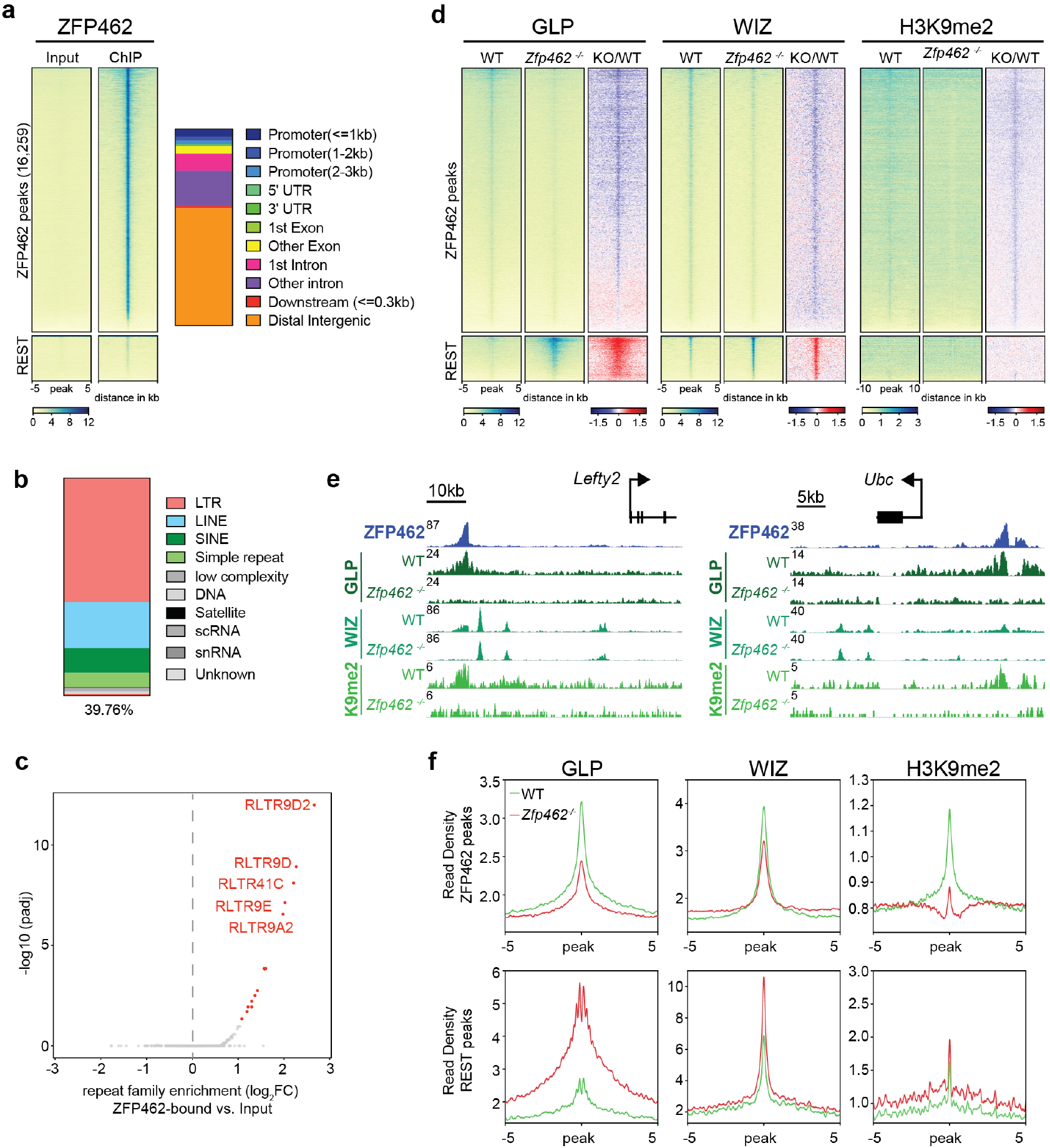
ZFP462 establishes H3K9me2 containing heterochromatin by recruiting GLP and WIZ proteins. **a)** Heatmap of ZFP462 ChIP–seq enrichment at significant peaks (16,259) in mESCs. Each row represents a 10 kb window centred on ZFP462 peak midpoint, sorted by ZFP462 ChIP signal. Input signals for the same window are shown on the left. Below are scale bars. (n = average distribution of two replicates). Bar plot, on the right, shows percentage of genomic features at ZFP462 ChIP-seq peaks. **b)** Bar plot shows fraction of repetitive DNA types at ZFP462 ChIP-seq peaks. 39.76 % of ZFP462 peaks are associated with repeat elements. **c)** Volcano plot shows enrichment and corresponding significance of TE families overlapping with ZFP462 peaks compared ChIP-seq input DNA. **d)** Heatmaps of GLP, WIZ and H3K9me2 ChIP–seq enrichment at ZFP462 and REST peaks in WT and *Zfp462^-/-^* mESCs. Blue to red scaled heatmap represents corresponding ChIP-seq enrichment ratio between *Zfp462^-/-^* versus WT (KO/WT). ChIP-seq signals are sorted by GLP enrichment ratio. For GLP and WIZ, each row represents a 10 kb window centred on ZFP462 peak midpoints. For H3K9me2, each row represents a 20 kb window (n = average distribution of two replicates). **e)** Screen shots of two selected genomic regions with ZFP462 peaks display GLP, WIZ and H3K9me2 ChIP-seq signals in WT and *Zfp462^-/-^* mESCs. ChIP-seq profiles are normalized for library size. **f)** Metaplots show average GLP, WIZ and H3K9me2 signal at ZFP462 and REST peaks in WT and *Zfp462^-/-^* mESCs. For each plot, read density is plotted at 10 kb window centred on peak midpoints.

To determine its in targeting the G9A/GLP complex to chromatin, we used CRISPR-mediated deletion to remove ZFP462 in Avi-GLP expressing mESCs and compared the genomewide enrichment of GLP to *Zfp462* wildtype cells (Extended Data Fig. 6b, c). In addition, we profiled WIZ and H3K9me2 in WT and *Zfp462* KO mESCs (Extended Data Fig. 6d). In wildtype mESCs, ZFP462 targets were co-occupied by GLP and WIZ, similar to REST binding sites, and displayed modest levels of H3K9me2 (Fig. 5d-f). In mESCs lacking ZFP462, enrichments of HMTase subunits and the histone methylation were markedly reduced at ZFP462 targets. By comparison, REST TF binding sites displayed increased GLP and WIZ ChIP-seq signals upon *Zfp462* deletion. Since global G9A levels remained unchanged in the mutant (Extended Data Fig. 5a), we conclude that the gain at REST binding sites reflects genomic redistribution of the H3K9-specific HMTase in the absence of ZFP462 interaction. Next, we used CRISPR to delete ZFP462 in Avi-HP1 expressing mESCs and tested its role in HP1 chromatin targeting (Extended Data Fig. 6c). Compared to known binding sites such as ADNP targets ^49^, Avi-HP1 showed only modest enrichment at ZFP462 peaks (Extended Data Fig. 6e). Moreover, the low HP1 signal remained largely unchanged by *Zfp462* deletion arguing against a significant role of ZFP462 in directly recruiting HP1 to its target loci. These results demonstrate that ZFP462 is primarily required for sequence-specific targeting of the G9A/GLP complex to induce heterochromatin at its targets.

### ZFP462-dependent heterochromatin restricts TF binding to TE-derived enhancers

TEs contribute up to 25% of binding sites for pluripotency TFs and play an integral role in wiring genes into the core regulatory network of mESCs^50,51^. Indeed, the mouse-specific RLTR9 subfamilies contain sequence motifs for pluripotency TFs including OCT4 and SOX2 and promote gene expression in mESCs, indicating bona fide enhancer activities^52–54^. Based on these studies, we performed H3K27ac ChIP-seq and ATAC-seq which confirmed that the vast majority of ZFP462 peaks also bear hallmarks of enhancer activity in mESCs (Fig. 6a-b, Extended Data Fig. 7a). Strikingly, DNA accessibility was specifically increased at ZFP462 target loci in KO mESCs while REST binding sites remained unchanged (Fig. 6a-c). This suggests that G9A/ GLP-dependent heterochromatin modifications restrict DNA accessibility potentially affecting TF occupancy at TE-derived enhancers. A search for known TF binding motifs at ZFP462 targets with increased DNA accessibility revealed highest enrichment of consensus sequences for core pluripotency TFs OCT4, SOX2, TCF and NANOG (Fig. 6d). Subsequent ChIP-seq profiling confirmed robust binding of OCT4, SOX2 and NANOG at ZFP462 target sequences in wildtype mESCs (Fig. 6e, Extended Data Fig. 7b-c). Notably, ZFP462 also colocalizes with these TFs at upstream regulatory sequences of the *Oct3/4* gene suggesting that it limits the expression of the pluripotency master regulator by targeting G9A/GLP (Extended Data Fig. 7b).

**Fig. 6:**
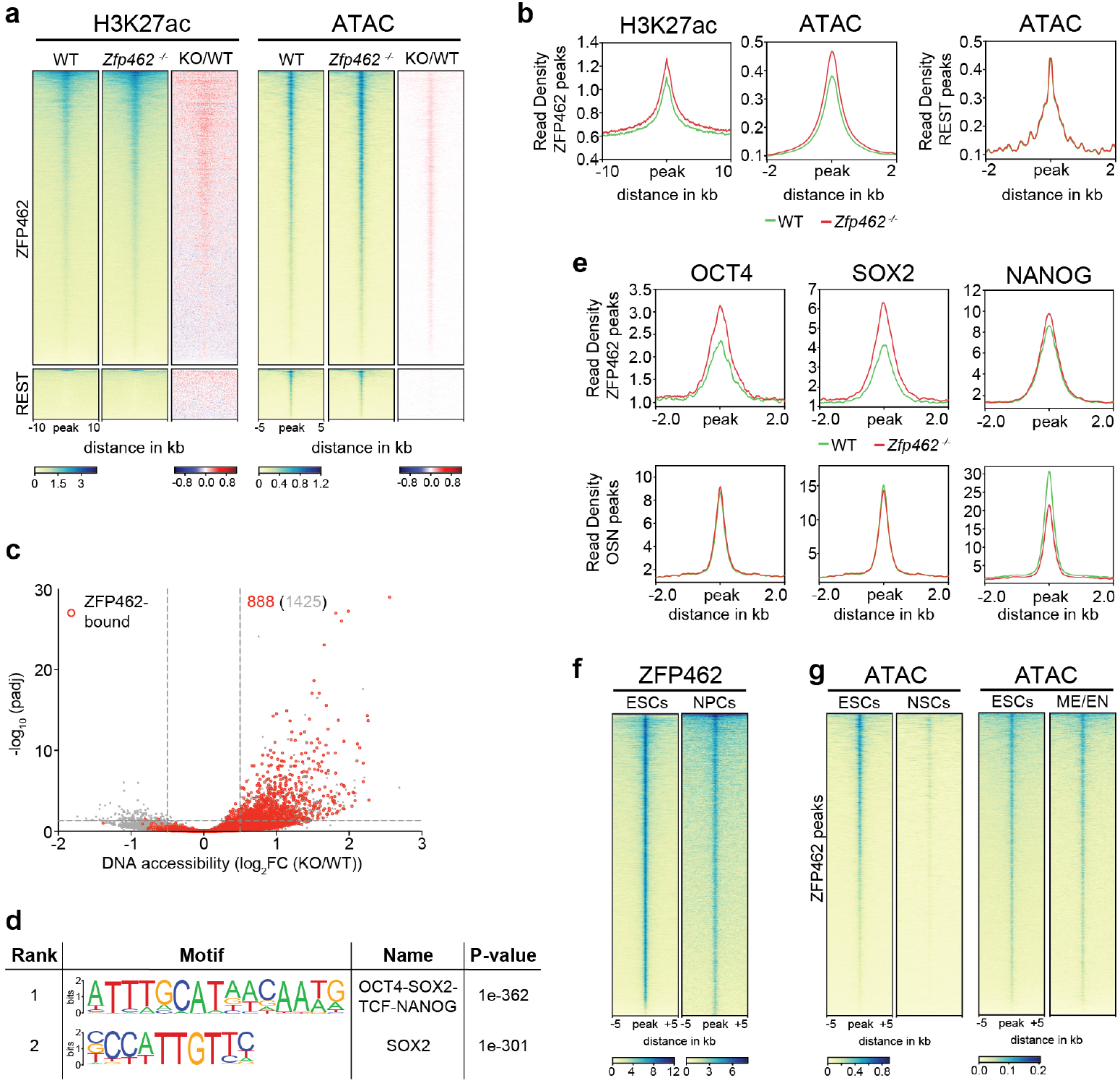
Zfp462 restricts binding of pluripotency transcription factors at lineage specifying enhancers protecting them from hyper-activation. **a)** Heatmaps of H3K27ac ChIP–seq and ATAC-seq at ZFP462 and REST peaks in WT and *Zfp462^-/-^* mESCs. Blue to red scaled heatmaps represent corresponding enrichment ratios between *Zfp462^-/-^* versus WT (KO/WT). H3K27ac rows represent a 20 kb window, ATAC-seq rows represent a 10 kb window centred on peak midpoints, sorted by H3K27ac enrichment ratio (n = average distribution of two H3K27ac ChIP-seq replicates and average distribution of three ATAC-seq replicates). **b)** Metaplots show average profiles of H3K27ac at ZFP462 peaks (left) and average profiles of ATAC-seq at ZFP462 and REST peaks (right). **c)** Volcano plot shows DNA accessibility changes and corresponding significance in WT and *Zfp462^-/-^* mESCs. Differentially accessible sites bound by ZFP462 are highlighted in red. Indicated are the number all loci with increased DNA accessibility and the number of ZFP462-bound loci with increased DNA accessibility. **d)** HOMER analysis of known DNA sequence motifs enriched at ZFP462-bound peaks with increased DNA accessibility. Top ranked DNA sequence motifs and respective significance values are shown in the table. **e)** Metaplots show average OCT4, SOX2 and NANOG ChIP-seq signal in WT and *Zfp462 ^-/-^* mESCs at ZFP462-bound peaks with increased DNA accessibility (KO/WT, padj. 0.05) and OCT4, SOX2 and NANOG overlapping peaks that are not ZFP462-bound in WT (OSN peaks). (n = average distribution of two replicates). **f)** Heatmap of ZFP462 ChIP-seq enrichment at ZFP462 peaks in mESCs and NPCs. ChIP-seq rows represent 10 kb window centred on ZFP462 ChIP–seq peak midpoints, sorted by ZFP462 ChIP-seq signal intensity (n = average distribution of two replicates). **g)** Heatmaps of ATAC-seq signal at ZFP462 peaks in mESCs and NSCs (left). Heatmaps of ATAC-seq signal at ZFP462 peaks in mESCs and mes-endoderm (ME/EN) cells ^60^. ATAC-seq rows represent 10 kb window centred on ZFP462 peak midpoints, sorted by mESCs ATAC-seq signal intensity.

The precise regulation of pluripotency TFs is essential for embryonic stem cell maintenance^55^. For example, a less than twofold increase in OCT4 expression is sufficient to trigger spontaneous differentiation of mESCs into primitive endoderm and mesoderm ^35,56–59^. To investigate if ZFP462 is required for restricting sequence interactions at TE-derived enhancers, we compared pluripotency TF binding at canonical OCT4-SOX2-NANOG targets (OSN peaks) and ZFP462 targets with significantly increased DNA accessibility in KO mESCs (padj. = 0.05) (Fig.6e, Extended Data Fig. 7c). Relative to wildtype mESCs OCT4 and SOX2 were markedly increased at *Zfp462* KO-sensitive enhancers. This change was specific since TF enrichments were unaffected at OSN peaks that do not overlap with ZFP462. In agreement with spontaneous differentiation into primitive endoderm, NANOG expression was downregulated resulting in reduced occupancy at OSN peaks. In contrast, its binding remained high at *Zfp462* KO-sensitive TE-derived enhancers, consistent with a relative increase in DNA accessibility.

Next, we interrogated our transcriptome data to test if increased OCT4 and SOX2 binding at ZFP462 bound TE-derived enhancers impacts transcriptional regulation. Genes proximal to ZFP462 targets with significantly increased DNA accessibility were more frequently upregulated arguing that enhanced TF binding promotes gene expression (Extended Data Fig. 8a). To test if proximal gene regulation is linked to spontaneous endodermal differentiation, we performed GO term analysis. Notably, the most enriched GO terms were related to developmental processes, in particular cardiovascular system development (Extended Data Fig. 8b). Finally, we analyzed TF binding motifs at ZFP462 targets with the highest increase in DNA accessibility (padj. = 0.05) and found that consensus sequences for SOX family TFs, including SOX17, were enriched (Extended Data Fig. 8c). Although SOX17 is only present at low levels in mESCs, increased accessibility of its binding motif might potentiate TF interaction frequency and ultimately promote upregulation of proximal genes associated with endodermal lineage specification. Together, these results demonstrate that ZFP462 binds to recently evolved, mouse-specific TEs and controls their regulatory activity in mESCs by targeting heterochromatin to restrict binding of pluripotency TFs including OCT4 and SOX2. In the absence of ZFP462, DNA accessibility is increased at TE-derived enhancers leading to increased expression of proximal genes involved in endoderm differentiation.

### Endodermal enhancers are ZFP462-bound and repressed in neural cells

Our data argue that ZFP462 is required for normal neuro-differentiation and development through regulation of TE-derived enhancers in neural cell lineages. To determine if ZFP462 occupies TE-derived enhancers later in development, we differentiated Avi-ZFP462-expressing mESCs and mapped its genome-wide distribution in NPCs. ZFP462 occupancy was largely conserved between mESCs and NPCs, arguing that this factor also prevents aberrant TF binding in neural cells (Fig. 6f). To investigate DNA accessibility at ZFP462 targets in neural cells, we used ATAC-seq in cultured neural stem cells (NSCs) originating from the subventricular zone of adult mice which showed robust ZFP462 protein expression (Extended Data Fig. 8d). This analysis revealed that ZFP462-bound enhancers are mostly inaccessible in wildtype NSCs and argues that G9A/GLP-dependent heterochromatin efficiently restricts inappropriate TF binding in neural cells (Fig. 6g).

Finally, we interrogated public ATAC-seq data to compare DNA accessibility at ZFP462 targets in endoderm progenitors^60^. In contrast to NSCs where ZFP462-bound enhancers are silent, endoderm progenitors displayed substantial DNA accessibility at these loci indicative of lineagespecific enhancer activity (Fig. 6g). Together, these results suggest that ZFP462 controls endoderm-specific enhancers by recruiting G9A/ GLP and establishing heterochromatin in mESCs and cells of the neural lineage. Our data suggest that by preventing lineage non-specific gene expression ZFP462 plays a central role in promoting robust differentiation along the neural lineage.

## Discussion

Using a CRISPR genetic screen for regulators of heritable H3K9me2/3-dependent gene silencing in mouse embryonic stem cells (mESCs), we discovered *Zfp462*, an essential gene encoding a C2H2 zinc-finger transcription factor, previously of unknown function. Heterozygous mutations of the human ortholog, *ZNF462*, cause Weiss-Kruszka syndrome, a genetic disorder characterized by a spectrum of neurodevelopmental defects including developmental delay, craniofacial abnormalities and autism^32^, but the underlying molecular mechanism is unknown. Our experiments demonstrate that ZFP462 is critical for the regulation of cell identity by restricting the unscheduled expression of endoderm-specific genes in mESCs and during neural differentiation.

ZFP462 binds to TE-derived enhancer sequences that contain ESC- and endoderm-specific TF binding motifs. Silencing of these loci is achieved through recruitment of G9A/GLP catalyzing H3K9me2. The establishment of heterochromatin at ZFP462 target sites restricts DNA accessibility and thereby precisely controls TF binding and enhancer activity. Failure to target G9A/GLP-dependent heterochromatin in heterozygous and homozygous *Zfp462* deletion mutants results in aberrant expression of endoderm-specifying transcription factors which impairs self-renewal in mESCs and interferes with timely establishment of neural-specific gene expression patterns during NPC differentiation. Neural differentiation is similarly impaired in heterozygous and homozygous Zfp462 mutant cells, consistent with the neurodevelopmental defects observed in patients with *ZNF462* haploinsufficiency.

Despite G9A/GLP - dependent heterochromatin modifications, ZFP462 target loci are co-occupied by core pluripotency TFs and show hallmarks of enhancer activity in mESCs. Transient co-localization of H3K9me2 and H3K27ac has also been reported at developmental enhancers which become decommissioned during *in vitro* differentiation of ESCs toward EpiSCs^61^. Thus, unlike KZFPs which enforce robust, transcriptional repression through Setdb1-dependent H3K9me3^62^, ZFP462 targets G9A/GLP to mediate a less restrictive heterochromatin that is compatible with DNA-protein interaction. Previous studies have shown that precise levels of OCT4 expression are critical for mESC self-renewal and govern cell type specification during early embryogenesis^35^. A less than two-fold increase in OCT4 expression in mESCs causes differentiation into primitive endoderm and mesoderm whereas reduction triggers differentiation into trophectoderm. Here we show that ZFP462 functions to limit DNA accessibility to impede OCT4 binding at TE-embedded enhancers preventing aberrant activation of an endoderm-specific gene regulatory network. Thus, by modulating OCT4 activity at ESC-specific enhancers, ZF462 may contribute to the quantitative readout of the master transcriptional regulator defining one of the earliest steps of cell fate specification during embryonic development.

Although ZFP462 is highly conserved across vertebrates, we found that in mESCs and NPCs it predominantly binds to evolutionarily young, species-specific TEs which entered the mouse genome <12-14 million years ago after the mouse-rat split^52^. How can this discrepancy be explained? One possibility is that ZFP462-bound TEs contain DNA sequence motifs that are much “older” that the rest of the TE sequence. Indeed, mouse-specific RLTR9 subfamilies harbor conserved binding sites for key pluripotency TFs including OCT4 and SOX2^52,53^. Adaptation of pluripotency TF binding sites bestows TEs with enhancer activity specific for germ cells and preimplantation embryos optimizing the likelihood to persist by vertical transmission^51,63,64^. We surmise that ZFP462/ZNF462 emerged in vertebrates to repress TEs which exploited this window of opportunity by recognizing a DNA sequence that overlaps with pluripotency TF binding motifs. Furthermore, by deploying G9A/ GLP to establish a less repressive heterochromatin domain ZFP462 may overcome the challenge to discriminate silencing of a mutagenic TEs from bona fide host gene regulatory sequences. As TE-derived enhancers have been co-opted to contribute to the ESC-gene regulatory network, so was the function of ZFP462 to limit DNA accessibility in order to precisely control the levels of pluripotency TF activity critical for cell specification.

Based on our findings in murine cells, we propose that the human ortholog ZNF462 plays a critical role in G9A/GLP regulation, targeting non-neural genes for repression during neurogenesis. Failure to restrict the expression of lineage nonspecific genes due to a ZNF462 heterozygous LOF mutation might impair proper neural cell differentiation and potentially underlie neurodevelopmental pathology. However, given the species-specificity of ZFP462-regulated TEs in mouse, we predict that ZNF462 will have novel human targets controlling a distinct gene regulatory network compared to mice. Exploring the human-specific functions of ZNF462 thus demands further investigation. Nevertheless, despite species-specific differences in target loci, our findings predict that the molecular mechanism of *ZNF462*-linked pathology involves loss of heterochromatin silencing at TE-derived enhancers and interference in neural differentiation by lineage non-specific gene expression. Our results argue that deregulation of neural lineage-specific gene expression is initiated at early steps of embryonic development consistent with the broad defects affecting cells of neuronal and neural crest origin. Together, this allows hypotheses on the etiology of Weiss-Kruszka syndrome to be developed and tested in the future.

## Acknowledgements

We are grateful to all members of the Bell and Brennecke laboratories for support, feedback and discussions. HP1 and GLP Avi-tagged cell lines are kind gifts from Marc Bühler and Joerg Betschinger labs at FMI, Switzerland. We thank Noelia Urban for providing mouse NSCs. We thank Christa Buecker, Fabio Mohn and Kristoffer Jensen for experimental advice and helpful discussions. We thank Frederic Berger, Luisa Cochella and Jakob Schnabl for manuscript reading and commenting on it. We thank Vienna Biocenter Core Facility Next Generation Sequencing. The GMI/IMBA/IMP Scientific Service units, especially the BioOptics facility and the Mass Spectrometry unit (Karl Mechtler & team) provided outstanding support. DNA methylation analysis by LC-MS/MS was performed by the Metabolomics Facility at Vienna BioCenter Core Facilities (VBCF), funded by the City of Vienna through the Vienna Business Agency. We thank Life Science Editors for editorial assistance. We apologize to colleagues whose work could not be cited due to space limitations. Funding: O.B., J.B., U.E. and S.M. were supported by the Austrian Academy of Sciences. O.B. was supported by the New Frontiers Group of the Austrian Academy of Sciences (NFG-05), the Human Frontiers Science Programme Career Development Award (CDA00036/2014-C), and start-up funding from the Norris Comprehensive Cancer Center at Keck School of Medicine of USC. R.Y. was supported by EMBO Long-Term Fellowship (ALTF 256-2015). D.S. acknowledges support from the Novartis Research Foundation and the European Research Council under the European Union’s (EU) Horizon 2020 research and innovation program grant agreement (ReadMe-667951).

## Author contributions

R.Y., K.S. and O.B. initiated and designed the study. R.Y and K.S generated cell lines, performed CRISPR-Cas9 genetic screen and differentiation assays. R.Y. performed all molecular biology experiments. P.H. carried out immuno-histochemistry. R.Y. and M.N analysed transcriptome and epigenome data. J.W. and G.M. analysed CRISPR-Cas9 genetic screen data. U.E., G.M and G.V. provided the CRISPR sgRNA library and helped with the screen design. L.M. and C.P. participated in experiments. L.I. and D.S. shared reagents and experimental expertise. J.B. co-supervised part of the project in his laboratory. O.B. supervised all aspects of the project. The manuscript was prepared by R.Y. and O.B.. D.S. and J.B. edited the manuscript. All authors discussed results and commented on the manuscript.

## Data availability

RNA-seq, QuantSeq, ATAC-seq and ChIP-seq data produced in this study are deposited at the Gene Expression Omnibus (GEO) database under super series accession number GSEXXXXXX. REST and ADNP ChIP-seq data are obtained from previously published datasets GSE27148^65^ and GSE97945^49^ respectively. Endodermal cells ATAC-seq data is obtained from previously published dataset GSE116262^60^.

## Materials and Methods

### Cell culture

Mouse Embryonic Stem Cells (mESCs) (TC1/ RRID:CVCL_M350) were cultured on gelatin-coated dishes in mESC medium containing DMEM supplemented with 15% fetal bovine serum (FBS; F7524/Sigma), 1× non-essential amino acids (Gibco), 1 mM sodium pyruvate (Gibco), 2 mM L-glutamine (Gibco), 0.1 mM 2-mercaptoethanol (Sigma), 50 mg ml^-1^ penicillin, 80 mg ml^-1^streptomycin and homemade LIF, at 37 °C in 5% CO2. Cells were tested for mycoplasma contamination every second week. Neural Stem Cells (NSCs) are isolated from subventricular zone of 7-8 week mouse brain and cultured as previously described ^66^.

### Generation of CiA Oct4 dual reporter mESCs

To generate CiA Oct4 dual reporter mESCs with TetO binding sites, first excised floxed Neomycin resistance cassette from Oct4 (CiA:Oct4) mESCs ^12^ by transfecting Cre-mCherry plasmid using Mouse Embryonic Stem Cell Nucleofector™ Kit from Lonza. After 24 hours after transfection, Cre-mCherry expressing cells were FACS sorted and sparsely seeded for clonal expansion on 15cm plate. The resulting clones were individually picked, split and screened by neomycin selection and genotyping for neomycin resistance cassette excised cells. Into these neomycin sensitive cells, TetO-BFP reporter cassette (7xTetO-PGK-Puro-BFP-pA) was introduced downstream of Oct4-GFP reporter using CRISPR/ Cas9 assisted homologous recombination by transfecting sgRNA/CAS9 ribonucleoprotein (RNP) complex and donor plasmid YR9. sgRNA and donor cassette sequences can be found in supplementary table 5. Stable Oct4 reporter cell line expressing GFP and BFP was isolated by FACS analysis and genotyping.

### TetR-FLAG-fusion construct design and delivery

All TetR-FLAG fusion constructs for recruitment at reporter locus were created as lentiviral plasmids expressing the gene of interest linked to TetR-3XFLAG-Gene-P2A-mCherry coding sequence under the control of an EF1a promoter. Lentivirus was produced by polyethylenimine (PEI) co-transfection of the desired construct and two packaging vectors VSV-G (Addgene #8454) and psPAX2 (Addgene #12260) in Lenti X 293T cells (Takara #632180). After 48–72 h, the virus was collected. Oct4 dual reporter cells were then transduced with the virus for 48 h in the presence of 8 μg/ml polybrene (Santa Cruz Biotechnology, SACSC-134220). Transduced cells with desired TetR-3XFLAG fusion constructs were isolated by FACS sorting for mCherry and western blot confirmation. Reversal of TetR fusion protein binding was achieved by addition of 1 μg/ml Doxycycline (final concentration / Sigma-D9891) to mESC culture medium.

### Genetic screen using CRISPR-Cas9 mutagenesis

For genetic screen, stable Cas9 expressing Oct4 dual reporter mESC line was generated by introducing Cas9-Blast through viral transduction into Oct4 dual reporter mESCs. After completion of blast selection, cells are sparsely seeded for clonal expansion on 15cm plate. The resulting clones were individually picked, split and screened for blast resistance. Into a blast resistant clone, pEF1a-TetR-3XFLAG-HP1-P2A-mCherry containing plasmid was introduced by transduction. mCherry expressing cells were FACS sorted and sparsely seeded for clonal expansion on 15cm plate. The resulting clones were individually picked, split and screened for initiation of reporter silencing and then for maintenance of reporter silencing by addition of doxycycline. From this selection process, mESC clone #H5 with Cas9-Blast / pEF1a-TetR-3XFlag-HP1-P2A-mCherry was selected for genetic screen.

For CRISPR/Cas9 mutagenesis, sgRNA library targeting nuclear factors described in^36^ was used. For retroviral library generation, barcoded sgRNA plasmid library carrying a neomycin-resistance cassette was packaged in PlatinumE cells (Cell Biolabs) according to the manufacturer’s recommendations. 300 million #H5 clone mESCs were infected with a 1:10 dilution of virus-containing supernatant for 24 h in the presence of 2 μg/ml polybrene (Santa Cruz Biotechnology, SACSC-134220) (Before infection, 300 million cells are divided into three sets, each set with 100 million cells. These three sets are treated as independent replicate till the completion of the screen and sequencing sgRNAs). Twenty-four hour after infection, selection for infected cells was started with G418 (Gibco) at 0.5 mg/ml. After twenty-four hour of selection, cells were split, and 480 million cells were seeded on sixty 15-cm dishes (Greiner; 639160, each replicate in 20 dishes). After that, cells were kept at a minimum number of 300 million cells (sum of three replicates) during editing and screening. After 5days of G418 selection, cells are cultured in doxycycline containing medium for 2 days.

Thereafter, GFP positive cells are FACS sorted. Unsorted mutant population was used as background control. From FACS sorted and unsorted cells, genomic DNA was isolated, sgRNA cassette was PCR amplified and sequenced on an Illumina HiSeq 2500, and data was analysed as previously described^36^. Gene enrichment was determined using MAGeCK^37^.

### Generation of *Zfp*462 endogenously tagged mESC line

For endogenous *Zfp462* tagging, 5 million Rosa26:BirA-V5-expressing cells^67^ were transfected with sgRNA/CAS9 ribonucleoprotein complex (sgRNA pre-incubated with CAS9 in cleavage buffer) mixed with 15ug of donor plasmid containing TAG(Avi-GFP-3XFLAG) sequence flanked on both sides by ~500bp of *Zfp462* start codon adjacent homology sequences. sgRNA and TAG sequence can be found in supplementary table 5. Transfection was carried out by electroporation using Mouse Embryonic Stem Cell Nucleofector™ Kit from Lonza. After 2 days of transfection GFP expressing cells were FACS sorted and sparsely seeded for clonal expansion on 15cm plate. The resulting clones were individually picked, split and screened by genotyping and western blot for desired tag integration.

### Generation of *Zfp462*^-/-^ and heterozygous mutant mESCs

*Zfp462^-/-^* mESCs were generated with CRISPR-Cas9 using two independent sgRNAs. Sequences of sgRNA can be found in supplementary table 5. The sgRNA sequences were cloned into the SpCas9-Thy1.1 plasmid. Cas9-sgRNA-Thy1.1 plasmid transfection was carried out by electroporation using Mouse Embryonic Stem Cell Nucleofector™ Kit from Lonza. After 36 hours of transfection Thy1.1 expressing cells were FACS sorted and sparsely seeded for clonal expansion on 15cm plate. The resulting clones were individually picked, split and screened by genotyping and western blot for *Zfp462^-/-^* mouse ESCs.

Heterozygous mutant mESCs are generated as previously described ^68^ with few modifications. For generation of *Zfp462^+/Y1195*^* and *Zfp462^+/R1257*^* heterozygous mutant mESCs, transfected sgRNA/ CAS9 ribonucleoprotein (RNP) complex and mix of 1:1 ratio WT/mutant donor single-stranded oligodeoxynucleotides carrying PAM mutation by electroporation using Mouse Embryonic Stem Cell Nucleofector™ Kit from Lonza. sgRNA and donor single-stranded oligodeoxynucleotide sequences can be found in supplementary table 5. One day after transfection, cells are sparsely seeded for clonal expansion on 15cm plate. The resulting clones were individually picked, split and screened by PCR followed by Restriction Fragment Length Polymorphism (RFLP). In case of *Zfp462^+/Y1195*^*, WT allele is resistant to MseI and mutant allele is sensitive to MseI. In case of *Zfp462^+/R1257*^*, WT allele is sensitive to TaqI and mutant allele is resistant to TaqI. Selected clones based on PCR-RFLP genotyping are further confirmed by sanger sequencing and western blot.

### Western blotting

Cells were grown to confluency on 10cm plates, collected in PBS, pelleted by 3 min centrifugation at 300*g*, and cell pellets were then resuspended in 5ml of buffer 1 (10 mM Tris-HCl at pH 7.5, 2 mM MgCl_2_, 3 mM CaCl_2_, Roche Complete protease inhibitor), incubated for 20 min at 4°C followed by a centrifugation step. The cell pellet was resuspended in buffer 2 (10 mM Tris-HCl at pH 7.5, 2 mM MgCl_2_, 3 mM CaCl_2_, 0.5% IGEPAL CA-630, 10% glycerol, Roche Complete Protease Inhibitor), incubated for 10 min at 4°C followed by centrifugation. Isolated nuclei were lysed in buffer 3 (50mM HEPES-KOH, pH7.3, 200mM KCl, 3.2mM MgCl_2_, 0.25% Triton X-100, 0.25% NP-40, 0.1% Na-deoxycholate and 1mM DTT and Roche Complete protease inhibitor), 2ul of Benzonase (E1014, Millipore) was added and incubated for 1 hour at 4°C. The lysate was cleared by centrifugation at 16,000*g* for 10 min at 4 °C, and the protein concentration in the nuclear extract was determined using the Broadford protein assay. For western blotting, 30 μg of protein was resolved on NuPAGE-Bis-Tris 4–12% mini protein gels (Invitrogen), which were transferred on to polyvinylidene fluoride (PVDF) membrane, blocked for 30 min in 5% non-fat dry milk in TBS with 0.1% Tween 20 (TBST), and stained with primary antibodies at 4 °C overnight. The primary antibodies used for western blotting were mouse anti-Flag (1:1,000, Sigma clone M2), rabbit anti-ZNF462(1:500, PA5-54585/Invitrogen), rabbit anti-Lamin B1(1:5,000, ab16048/abcam), rabbit anti-G9a (1:1,000, 68851S/Cell signaling) and mouse anti-H3K9me2(1:1,000, ab1220/abcam), Signal was detected with corresponding horseradish peroxidase (HRP)-conjugated secondary antibodies and Clarity Western ECL substrate (170-5061, Bio-Rad).

### Differentiation of mESCs to Neural Progenitor Cells (NPCs) and immunostaining

Differentiation was performed as previously described ^46^ with few modifications. mESCs are cultured on gelatin coated plates. In addition, cells are cultured in mESC medium containing 2i inhibitors (3 μM glycogen synthase kinase (GSK) inhibitor and 10 μM MEK inhibitor) for five passages before starting the differentiation experiment.

For immunostaining, four-day retinoic acid treated EBs were fixed with 4% PFA (Thermo Scientific, #28906) for 30 min, washed 3x with PBS and stored at 4 °C until further processing. EBs were incubated in blocking solution, which consisted of PBS (Gibco, #14190094), 4% goat/donkey serum (Bio-Rad Laboratories, #C07SA/Bio-Rad Laboratories, #C06SB) and 0.2% Triton X-100 (Sigma-Aldrich, #T8787) for at least 15 min before application of primary antibody in the same blocking solution. The primary antibody was incubated with the EBs for 2 days at 4C while shaking. Used primary antibodies were SOX1 (R&D Systems, #AF3369, 1:200) and FOXA2 (Cell Signaling Technologies, #8186T, 1:200). Subsequently, EBs were washed twice for 15 min each using a PBS/ Tween20 solution (0.1% Tween20, Sigma-Aldrich, #P1379). A secondary antibody was then applied 1:500 in blocking solution for another 2 days at 4C, while shaking. Secondary antibodies used were: Donkey anti-Goat IgG (H+L) Highly Cross-Adsorbed Secondary Antibody, Alexa Fluor Plus 488 (Invitrogen, #A32814) and Goat anti-Rabbit IgG (H+L) Highly Cross-Adsorbed Secondary Antibody, Alexa Fluor Plus 647 (Invitrogen, # A32733). DAPI (Sigma-Aldrich, #9542) was added to the secondary antibody staining solution. Afterward, EBs were washed once with PBS/Tween20 solution for 15 min followed by a second wash with the same solution for 24h at 4C while shaking. Finally, EBs were stored at 4C prior to imaging on glass slides, in FocusClear (CellExplorer Labs, #FC-101) clearing solution. Image acquisition was carried out using Spinning Disk Confocal Olympus.

### Transcriptome analysis by RT–qPCR, RNA-seq and QuantSeq

For RT-qPCR experiments, total RNA was extracted from cells using the Qiagen RNeasy mini kit with on-column DNase digestion step. Total RNA (500 ng) was reverse transcribed using the SuperScript™ III Reverse Transcriptase (Invitrogen). RT–qPCR was performed on a CFX96 Real-Time PCR System (Bio-Rad) using the Promega GoTaq^®^ qPCR Master Mix (Ref: A6001) with RT-qPCR primers described in supplementary table 5. Relative RNA levels were calculated from *C*_t_ values according to the Δ*C*_t_ method and normalized to *Tbp* mRNA levels. For RNA-seq, total RNA was subjected to polyA enrichment using NEBNext^®^ Poly(A) mRNA Magnetic Isolation Module followed by library construction using the NEBNext^®^ Ultra™ II RNA Library Prep Kit for Illumina. QuantSeq libraries are generated using QuantSeq 3’ mRNA-Seq Library Prep Kit FWD for Illumina (Lexogen) according to manufacturer instructions. RNA-seq libraries were sequenced on Illumina NovaSeq machine with 100bp single-end sequencing and QuantSeq libraries are sequenced on NextSeq550 machine with 75bp singleend sequencing.

### Chromatin immunoprecipitation (ChIP)

For Chromatin Immunoprecipitation, 25 × 10^6^ mES cells were collected, washed in once in 1× PBS and crosslinked with formaldehyde at a final concentration of 1% for 7 min. The crosslinking was stopped on ice and with glycine at final 0.125 M concentration. The crosslinked cells were pelleted by centrifugation for 5 min at 1200 × *g* at 4 °C. Nuclei were prepared by washes with NP-Rinse buffer 1 (final: 10 mM Tris pH 8.0, 10 mM EDTA pH 8.0, 0.5 mM EGTA, 0.25% Triton X-100) followed by NP-Rinse buffer 2 (final: 10 mM Tris pH 8.0, 1 mM EDTA, 0.5 mM EGTA, 200 mM NaCl). Afterwards the cells were prepared for shearing by sonication by two washes with Covaris shearing buffer (final: 1 mM EDTA pH 8.0, 10 mM Tris-HCl pH 8.0, 0.1% SDS) and resuspension of the nuclei in 0.9 mL Covaris shearing buffer (with 1 × protease inhibitors complete mini (Roche)). The nuclei were sonicated for 15 min (Duty factor 5.0; PIP 140.0; Cycles per Burst 200; at 4 °C) in 1 ml Covaris glass cap tubes using a Covaris E220 High Performance Focused Ultrasonicator. Input samples were prepared from 25 μL of sonicated lysate. For this, chromatin was treated with RNase A and Proteinase K and crosslink reversed overnight at 65 °C. Sample is phenol chloroform extracted and DNA was precipitated. Shearing of DNA was confirmed to be between 250bp and 800 bp by agarose gel electrophoresis, also DNA concentration was estimated. Crude chromatin lysate was clarified by spinning at 20,000 × *g* at 4 °C for 15 min to separate insoluble debris. In parallel, human Embryonic Stem Cells (hESCs) chromatin was also prepared as mentioned above. After quantification, mESCs chromatin was spiked in with 1% of hESCs chromatin. Spiked in clear chromatin is used for immunoprecipitation. Chromatin equivalent of 50 μg DNA was incubated overnight in 1× IP buffer (final: 50 mM HEPES/KOH pH 7.5, 140 mM NaCl, 1 mM EDTA, 1% Triton X-100, 0.1% DOC, 0.1% SDS) with 5ul of each antibody (mouse anti-H3K9me2/ab1220/abcam, rabbit anti-H3K27ac/ab4729/abcam, rabbit anti-WIZ/ NBP1-80586/Novus, rabbit anti-OCT4/ab19857/ abcam, goat anti-SOX2/AF2018 and rabbit anti-NANOG/ab80892/abcam) at 4 °C on a rotating wheel. The overnight IPs were incubated with BSA-blocked Protein G coupled Dynabeads (Thermo Fisher Scientific) for 3 hours at 4 °C on a rotating wheel. Beads were subsequently washed 2× with IP buffer (final: 50 mM HEPES/KOH pH 7.5, 140 mM NaCl, I mM EDTA, 1% Triton-X100, 0.1% DOC, 0.1% SDS), 1× with high salt buffer (final: 50 mM HEPES/KOH pH 7.5, 500 mM NaCl, 1mM EDTA, 1% Triton-X100, 0.1% DOC, 0.1% SDS), 2× with DOC buffer (10 mM Tris pH 8, 0.25 mM LiCl, 1 mM EDTA, 0.5% NP40, 0.5% DOC) and 1× with TE (+50 mM NaCl). The DNA was then eluted two times with 150 μL Elution buffer (final: 1% SDS, 0.1 M NaHCO3) for 20 min each at 65 °C. The eluate was treated with RNase A and Proteinase K and crosslink reversed overnight at 65 °C. Next day eluate was phenol/chloroform extracted and IP DNA is ethanol precipitated.

ZFP462 and HP1 biotin ChIP was performed as previously described^67^ and GLP ChIP was performed as described previously^69^.

### ChIP–qPCR analysis and ChIP–seq libraries preparation

ChIP enriched DNA was subjected to qPCR analysis on a CFX96 Real-Time PCR System (Bio-Rad) using the Promega GoTaq^®^ qPCR Master Mix (Ref: A6001) with ChIP-qPCR primers described in Supplementary table 5. Relative ChIP enrichment was calculated by percent input method. For ChIP–seq sample preparation, library construction was performed using the NEBNext Ultra-II kit (New England Biolabs) following manufacturer recommendations. Libraries were sequenced on Illumina NextSeq550 machines, with 75-bp single-end sequencing.

### Protein Co-Immunoprecipitation (Co-IP)

Cells were grown to confluency on 15cm plates, trypsinised, collected in PBS by and pelleted by 3 min centrifugation at 300*g*. For each Co-IP, 30 million equivalent cell pellets were then resuspended in 5ml of buffer 1 (10 mM Tris-HCl at pH 7.5, 2 mM MgCl_2_, 3 mM CaCl_2_, Roche Complete protease inhibitor), incubated for 20 min at 4°C and collected cells by a centrifugation step. The cell pellet was resuspended in buffer 2 (10 mM Tris-HCl at pH 7.5, 2 mM MgCl_2_, 3 mM CaCl_2_, 0.5% IGEPAL CA-630, 10% glycerol, Roche Complete Protease Inhibitor), incubated for 10 min at 4°C followed by centrifugation to collect nuclei. Isolated nuclei were lysed in buffer 3 (50mM HEPES-KOH, pH7.3, 200mM KCl, 3.2mM MgCl_2_, 0.25% Triton X-100, 0.25% NP-40, 0.1% Na-deoxycholate and 1mM DTT and Roche Complete protease inhibitor). To this 4ul of Benzonase (E1014, Millipore) was added and incubated for 1 hour at 4°C rotating. The lysate was cleared by centrifugation at 16,000*g* for 10 min at 4 °C. The lysate was cleared by centrifugation and incubated for 2 hours at 4°C with Tag specific magnetic beads (Dynabeads™ M-280 Streptavidin/Invitrogen for immunoprecipitation of Avitag-ZFP462 and Avitag-GLP. Anti-FLAG^®^ M2 magnetic beads/Sigma for immunoprecipitation of TetR-3xFLAG-CT-ZFP462). The beads were washed four times for 10 min with buffer 4 (50mM HEPES-KOH, pH7.3, 300mM KCl, 3.2mM MgCl_2_, 0.25% Triton X-100, 0.25% NP-40, 0.1% Na-deoxycholate and 1mM DTT). Beads are further washed four times with Tris buffer (20mM Tris pH7.5, 137mM Nacl) and subjected for mass spectrometry analysis.

### Identification of proteins by mass spectrometry

Beads were resuspended in 50ul of 100 mM ammonium bicarbonate (ABC), supplemented with 400ng of lysyl endopeptidase (Lys-C, Fujifilm Wako Pure Chemical Corporation) and incubated for 4 hours on a Thermo-shaker with 1200 rpm at 37°C. The supernatant was transferred to a fresh tube and reduced with 0.5mM Tris 2-carboxyethyl phosphine hydrochloride (TCEP, Sigma) for 30 minutes at 60°C and alkylated in 3 mM methyl methanethiosulfonate (MMTS, Fluka) for 30 min at room temp protected from light. Subsequently, the sample was digested with 400ng trypsin (Trypsin Gold, Promega) at 37°C overnight. The digest was acidified by addition of trifluoroacetic acid (TFA, Pierce) to 1%. A similar aliquot of each sample (30%) was analysed by LC-MS/ MS using NanoLC-MS analysis. The nano HPLC system used was an UltiMate 3000 RSLC nano system (Thermo Fisher Scientific, Amsterdam, Netherlands) coupled to a Q Exactive HF-X mass spectrometer (Thermo Fisher Scientific, Bremen, Germany), equipped with a Proxeon nanospray source (Thermo Fisher Scientific, Odense, Denmark). Peptides were loaded onto a trap column (Thermo Fisher Scientific, Amsterdam, Netherlands, PepMap C18, 5 mm × 300 μm ID, 5 μm particles, 100 Å pore size) at a flow rate of 25 μL min-1 using 0.1% TFA as mobile phase. After 10 min, the trap column was switched in line with the analytical column (Thermo Fisher Scientific, Amsterdam, Netherlands, PepMap C18, 500 mm × 75 μm ID, 2 μm, 100 Å). Peptides were eluted using a flow rate of 230 nl min-1, and a binary 3h gradient, respectively 225 min. The gradient starts with the mobile phases: 98% A (water/formic acid, 99.9/0.1, v/v) and 2% B (water/acetonitrile/formic acid, 19.92/80/0.08, v/v/v), increases to 35%B over the next 180 min, followed by a gradient in 5 min to 90%B, stays there for 5 min and decreases in 2 min back to the gradient 98%A and 2%B for equilibration at 30°C. The Q Exactive HF-X mass spectrometer was operated in data-dependent mode, using a full scan (m/z range 380-1500, nominal resolution of 60,000, target value 1E6) followed by 10 MS/MS scans of the 10 most abundant ions. MS/MS spectra were acquired using normalized collision energy of 28, isolation width of 1.0 m/z, resolution of 30.000 and the target value was set to 1E5. Precursor ions selected for fragmentation (exclude charge states unassigned, 1, 7, 8, >8) were put on a dynamic exclusion list for 60 s. Additionally, the minimum AGC target was set to 5E3 and intensity threshold was calculated to be 4.8E4. The peptide match feature was set to preferred and the exclude isotopes feature was enabled.

For peptide identification, the RAW-files were loaded into Proteome Discoverer (version 2.5.0.400, Thermo Scientific). All hereby created MS/MS spectra were searched using MSAmanda v2.0.0.16129, Engine version v2.0.0.16129 ^70^. For the 1st step search the RAW-files were searched against the database mouse_uniprot_reference_2020-12-05.fasta (21,966 sequences; 11,718,975 residues), supplemented with common contaminants, using the following search parameters: The peptide mass tolerance was set to ±5 ppm and the fragment mass tolerance to 6ppm. The maximal number of missed cleavages was set to 2, using tryptic enzymatic specificity. The result was filtered to 1 % FDR on protein level using Percolator algorithm ^71^ integrated in Thermo Proteome Discoverer. A sub-database was generated for further processing. For the 2nd step the RAW-files were searched against the created sub-database using the following search parameters: Beta-methylthiolation on cysteine was set as a fixed modification, oxidation on methionine and deamidation on asparagine and glutamine were set as variable modifications. Monoisotopic masses were searched within unrestricted protein masses for tryptic enzymatic specificity respectively. The peptide mass tolerance was set to ±5 ppm and the fragment mass tolerance to ±6 ppm. The maximal number of missed cleavages was set to 2. The result was filtered to 1 % FDR on protein level using Percolator algorithm. Peptide areas have been quantified using in-house-developed tool apQuant^72^. Proteins were quantified by summing unique and razor peptides. Protein-abundances-normalization was done using sum normalization. Statistical significance of differentially expressed proteins was determined using limma ^73^.

### ATAC-seq

ATAC-seq was performed on WT and *Zfp462* knockout mESCs, and WT mouse NSCs following a previously published protocol^74^. The experiment was performed in biological replicates. Libraries were paired-end sequenced (2 × 75 bp) using an Illumina NextSeq550 sequencer.

### DNA methylation analysis by mass spectrometry

1ug of pure genomic DNA was digested using DNA Degradase plus (Zymo Research) for 6 hours at 37 degrees. For determination of the methyl-dC content, samples were analyzed with LC-MS/MS using a triple quadrupole mass spectrometer, employing the selected reaction monitoring (SRM) mode of the instrument. The following transitions were used in the positive ion mode: dC *m/z* 228.1 → *m/z* 112.1 and methyl-dC *m/z*242.1 → *m/z* 126.1. Data were interpreted using the Trace Finder software suite (Thermo Fisher Scientific) and manually validated. Calibration curves of defined mixtures of dC and methyl-dC were acquired using the same procedure and employed for calculating the molar percentage of methyl-dC.

### Phylogenetic analysis

ZFP462 orthologs were collected in a convergent PSI-BLAST search against the NCBI non-redundant protein database using amino-acids 632-835 of NP_766455.2 as a query. The obtained sequence set was filtered, removing identical and partial sequences, keeping only sequences with identity below 95%. The remaining sequences were aligned using IQ-TREE ^75^ version 1.6.1 using parameters -alrt 1000 -bb 1000, and the phylogenetic tree was visualized using iTOL ^76^.

### RNA-seq and QuantSeq data analysis

RNA-seq reads were trimmed using Trim Galore v0.5.0, whereas QuantSeq reads are trimmed using BBDuk v38.06. Trimmed reads mapping to abundant sequences included in the iGenomes Ensembl GRCm38 bundle (mouse rDNA, mouse mitochondrial chromosome, phiX174 genome, adapter) were removed using bowtie2 v2.3.4.1 alignment^77^. Remaining reads were analyzed using genome and gene annotation for the GRCm38/mm10 assembly obtained from Mus musculus Ensembl release 94. Reads were aligned to the genome using STAR v2.6.0c and reads in genes were counted with featureCounts (subread v1.6.2) with parameter -s 2 for RNAseq and using strand-specific read counting for QuantSeq samples with parameter -s 1^78^. Differential gene expression analysis on raw counts and variance-stabilized transformation of count data for heatmap visualization were performed using DESeq2 v1.18.1^79^. Functional annotation enrichment analysis of differentially expressed genes was conducted using clusterProfiler v3.6.0 in R v3.4.1^80^.

### ChIP-seq data analysis

ChIP-seq reads were trimmed using Trim Galore v0.4.4 and thereafter aligned to the mm10 reference genome using BWA MEM v0.7.17. Duplicated reads were marked and excluded with Picard MarkDuplicates v2.23.4 and the obtained bam files were used to generate bigwig files using deepTools bamCoverage v3.5.0 with the parameter -normalizeUsing RPKM. Peaks were called from the sorted BAM using MACS2 v2.1.1 with default settings^81^. To analyze overlapping and non-overleaping peaks, bedtools intersect command was used. Peaks in blacklist regions were identified using bedtools intersect v2.27.1 and mm10.blacklist.bed.gz v1. Spike-in normalization of ChIP data was performed as previously described^82^. Repeat analysis was performed using RepEnrich2 following alignment of trimmed reads with bowtie2 v2.2.9. Heatmaps and metaplots are generated using deepTools version 3.1.2^83^.

### ATAC-seq data analysis

ATAC-seq reads were processed using nfcore/atacseq v1.0.0 with standard settings. Reads were trimmed using trim-galore v0.5.0, aligned to the mm10 genome using BWA MEM v0.7.17 and duplicated reads were marked with Picard MarkDuplicates v2.19.0. Alignment was filtered using samtools v1.9 thereby removing multimapping and duplicated reads. Peaks were called from the filtered and sorted BAM using MACS2 v2.1.2 with parameters --broad --BAMPE --keep-dup all --nomodel. Consensus peaks were obtained by merging peak calls using bedtools mergeBed v2.27.1. Heatmaps and metaplots are generated using deepTools version 3.1.2. DNA sequence motif analysis in ATAC-seq peaks was performed using HOMER version homer/4.10-foss-2018b^84^.

## Supplementary Figures

**Extended Data Fig. 1.**
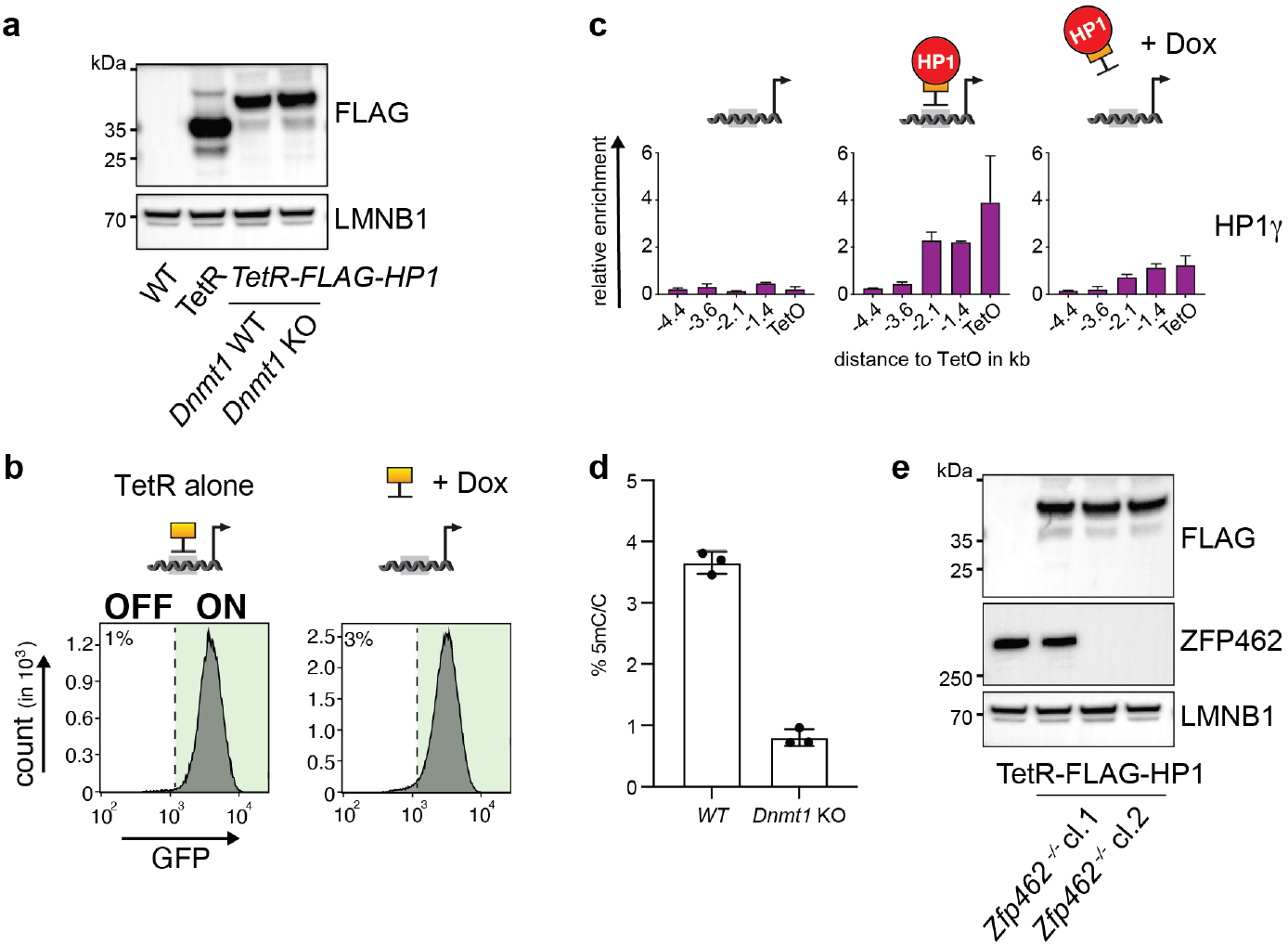
Characterization of WT and mutant CiA *Oct4* dual reporter cells. **a)** Western blot confirms expression of TetR-FLAG and TetR-FLAG-HP1 proteins in WT and *Dnmt1* knockout (KO) CiA *Oct4* dual reporter mESCs. LMNB1 is used as protein loading control. **b)** Flow cytometry histograms show GFP expression in CiA *Oct4* dual reporter cells after TetR-FLAG recruitment and after TetR-FLAG release following Dox addition for four days. **c)** ChIP-qPCR shows relative enrichment of HP1 surrounding TetO before TetR-FLAG-HP1 tethering, in the presence of TetR-FLAG-HP1 and after Dox-dependent release of TetR-FLAG-HP1 for four days. Data are mean ± SD (error bars) of at least two independent experiments. **d)** Bar plot shows fraction of cytosine methylation (5mC) in WT and *Dnmt1* KO CiA *Oct4* dual reporter mESCs measured by LC-MS. (n= three replicates) **e)** Western blot shows expression of TetR-Flag-HP1 and ZFP462 in WT and two independent *Zfp462* KO CiA *Oct4* dual reporter mESC lines. LMNB1 is used as loading control.

**Extended Data Fig. 2.**
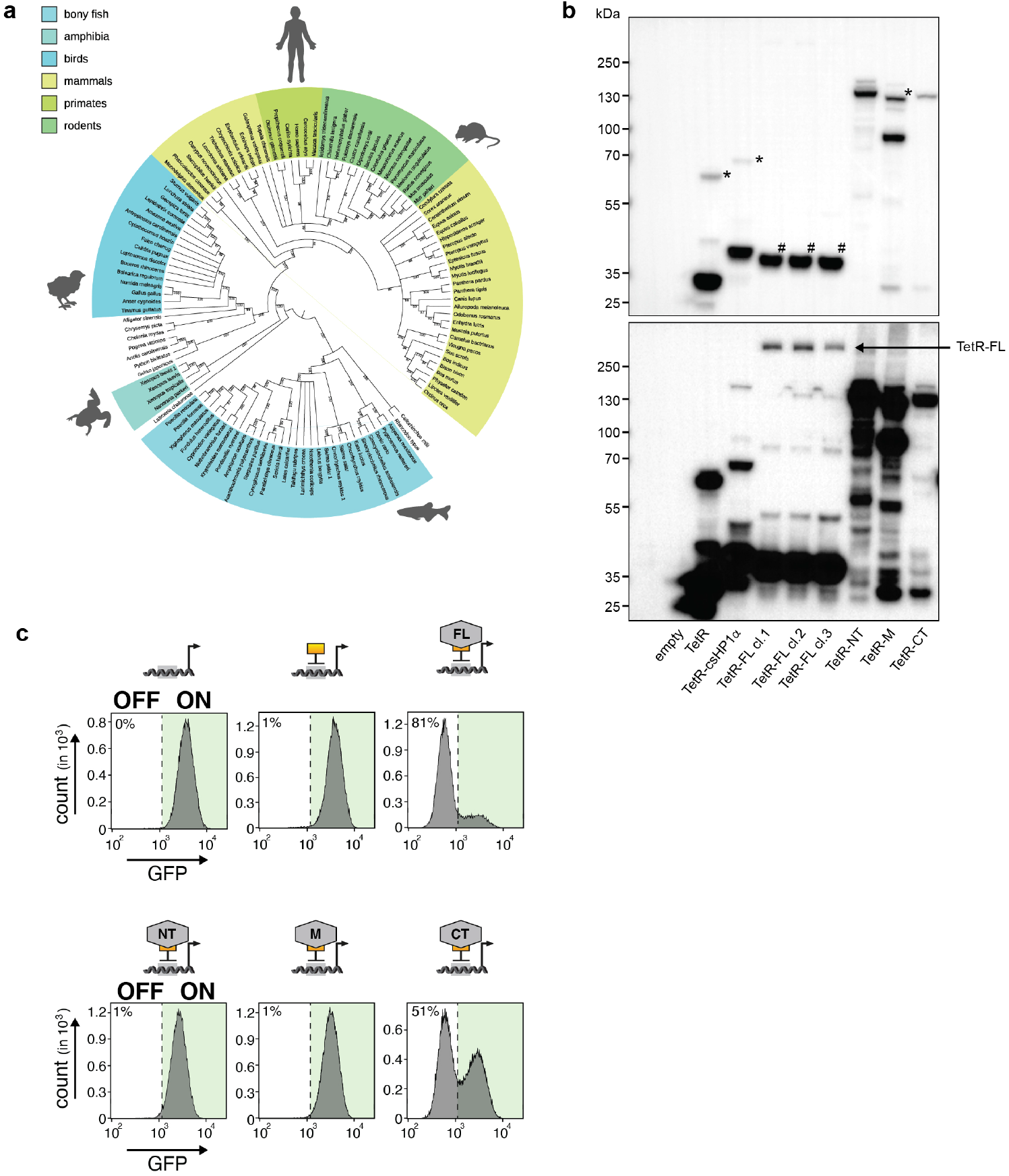
ZFP462 is conserved across vertebrates and acts as transcriptional repressor. **a)** Phylogenetic tree of ZFP462 protein orthologues in vertebrate species. Bootstrap values are shown on branches. **b)** Western blot shows expression ZFP462 fusions with TetR-FLAG in CiA *Oct4* dual reporter mESCs. Bottom picture is overexposed image. Hashtag symbol points cleaved TetR-FLAG protein from TetR-FLAG-ZFP462-FL (full Length). TetR-FLAG fusions initially express mCherry for selection which is cleaved via P2A signal. Asterisk marks P2A un-cleaved protein product. **c)** Representative flow cytometry histograms show GFP expression in CiA reporter mESCs expressing TetR-FLAG fusions with full-length or truncated ZFP462 protein. Each histogram is average profile of 100,000 analysed cells.

**Extended Data Fig. 3.**
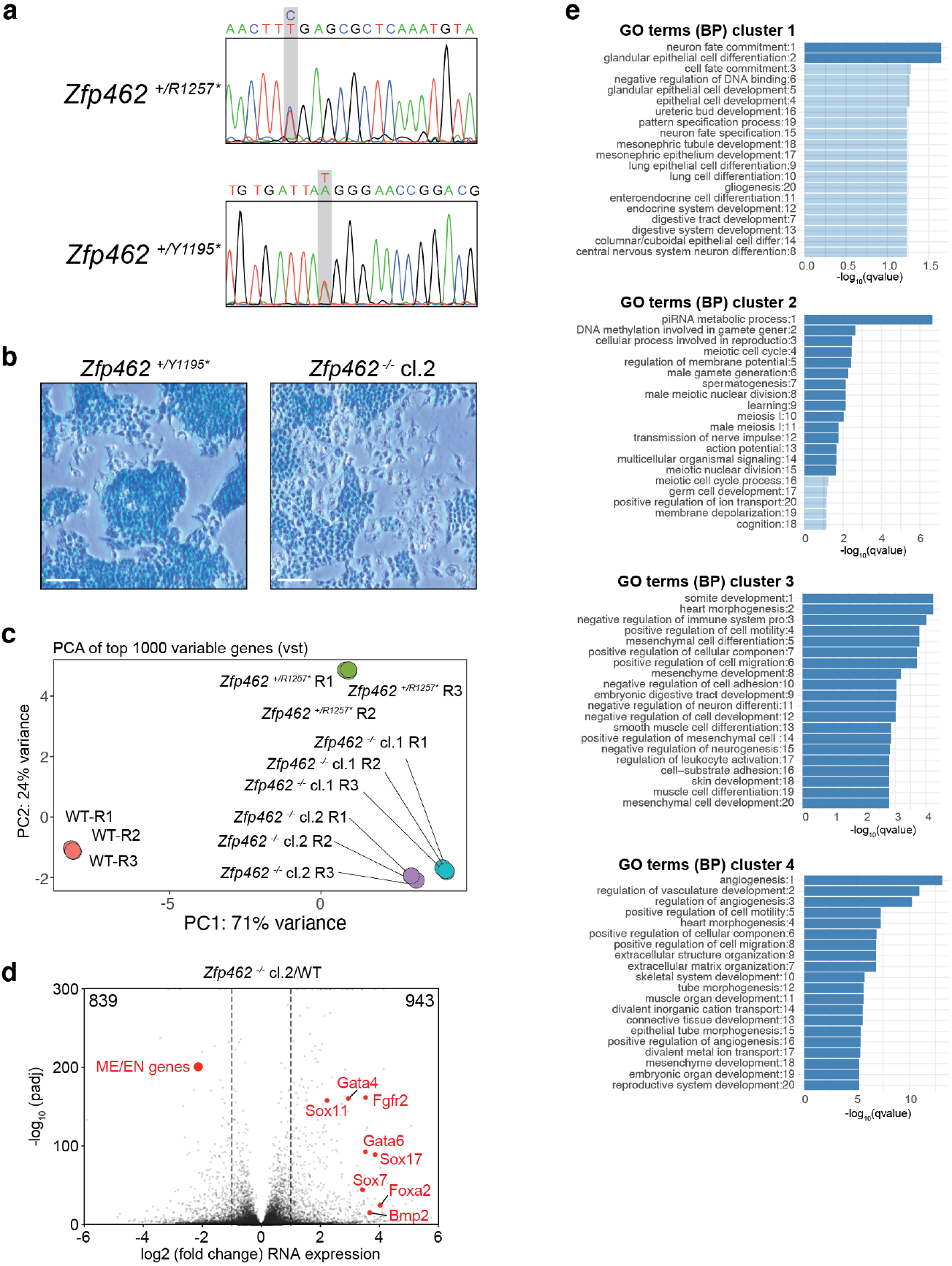
Meso-endodermal (ME/EN) genes are upregulated in *Zfp462* mutant mESCs. **a)** Sanger sequence chromatograms of heterozygous *Zfp462* mutants. Heterozygous non-sense mutations are highlighted in grey. **b)** Alkaline phosphatase staining of *Zfp462* ^+/Y1195*^ and *Zfp462* ^-/-^ *cl.2* mESCs (scale bar = 100 m). **c)** Correlation plot shows principal component analysis (PCA) of replicate RNA-seq experiments from WT, two *Zfp462 ^+/-^* and two *Zfp462 ^-/-^* mESC lines. **d)** Volcano plot shows differential gene expression of *Zfp462* ^-/-^ *cl.2* mESCs compared to WT mESCs. (n = three replicates). **e)** Bar plots show gene ontology (GO) terms enriched the four clusters of heatmap in Figure 3e.

**Extended Data Fig. 4.**
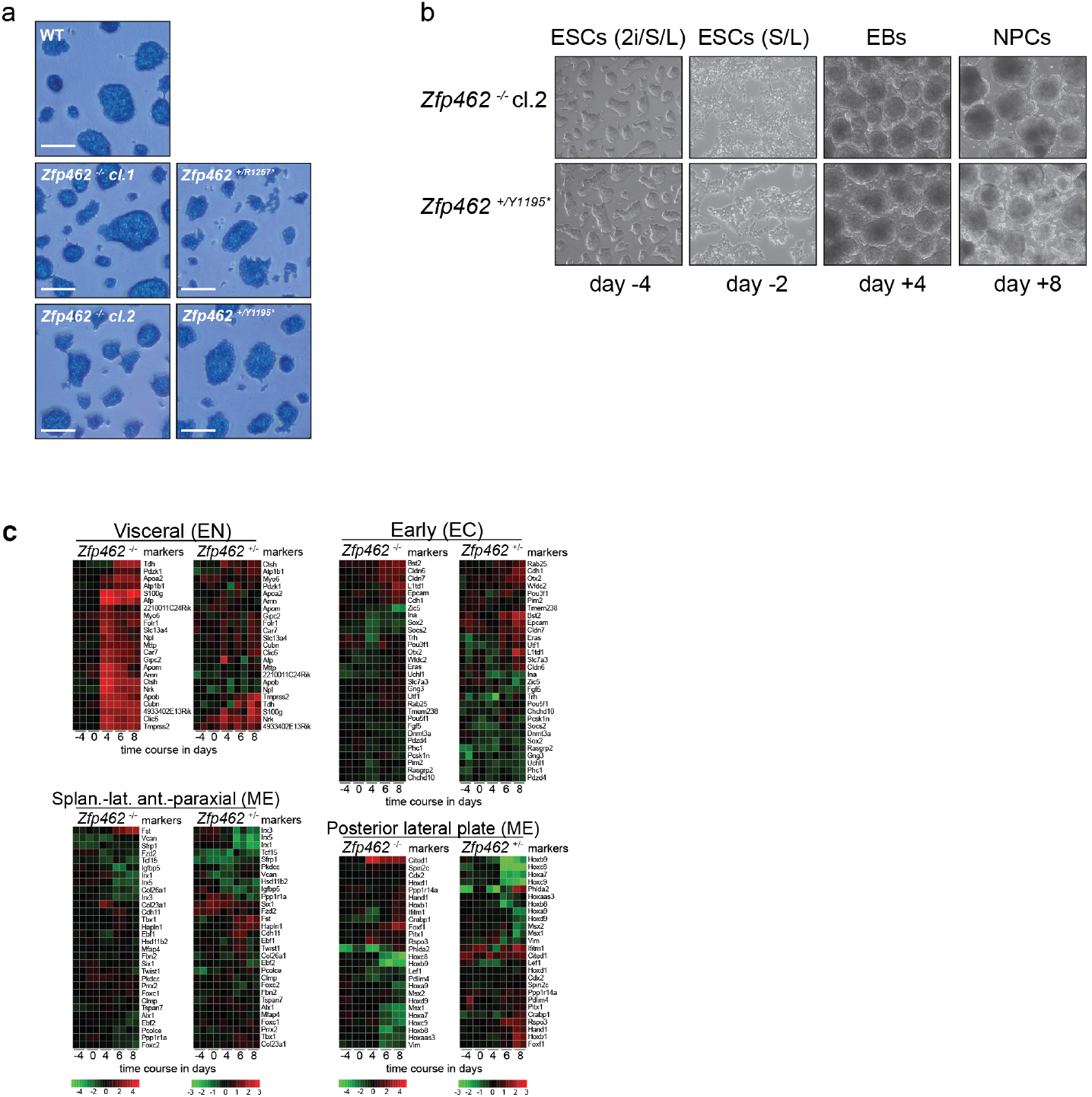
Lineage-specifying genes are deregulated during neuronal differentiation. **a)** Alkaline phosphatase staining of WT and *Zfp462* mutant mESCs cultured in 2i/LIF/Serum medium (scale bar = 180 μm). **b)** Representative bright field images show *Zfp462*^+/Y1195*^ and *Zfp462* ^-/-^ *cl.2* at corresponding stages of neural differentiation. **c)** Heatmaps show differential expression of selected marker genes specific for endodermal and neural lineages in heterozygous and homozygous *Zfp462* mutant cells during neural differentiation.

**Extended Data Fig. 5.**
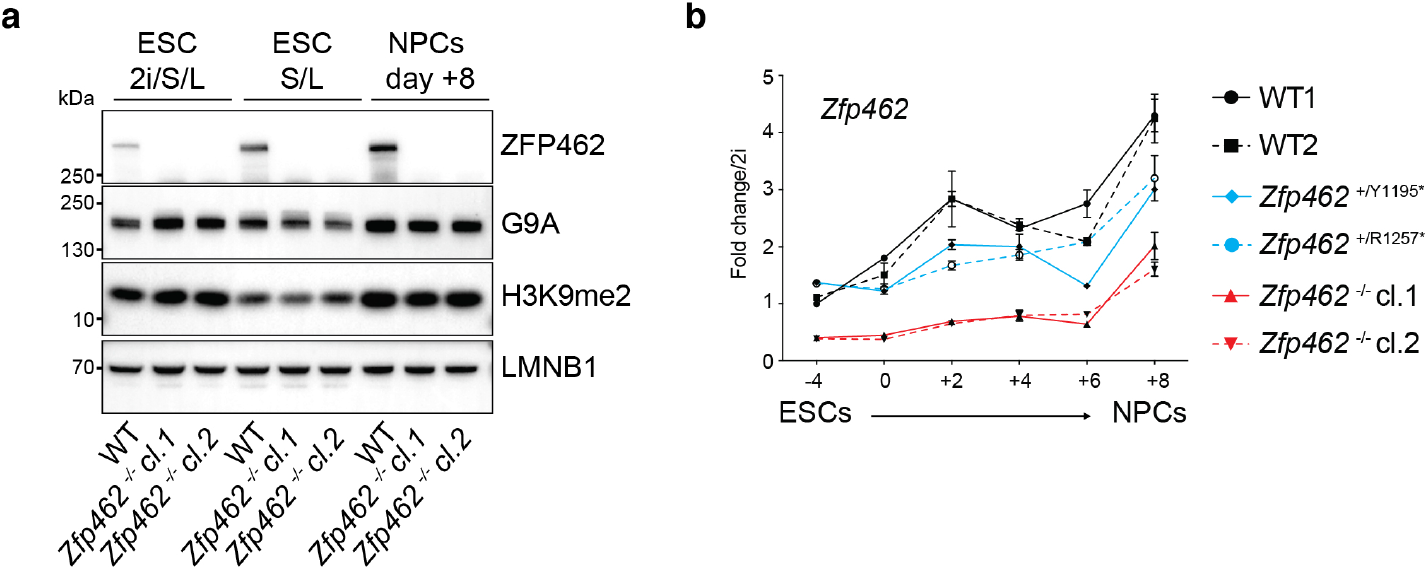
Analysis of ZFP462, G9A and H3K9me2 protein levels and *Zfp462* RNA expression level during neuronal differentiation. **a)** Western blot analysis shows levels of ZFP462, G9A and H3K9me2 in WT and *Zfp462* ^-/-^ mESCs during neural differentiation. LMNB1 is used as loading control. **b)** Line plot shows RT-qPCR analysis of *Zfp462* RNA expression during neural differentiation (n = two replicates). Expression level is shown relative to mESCs (2i/S/ L).

**Extended Data Fig. 6.**
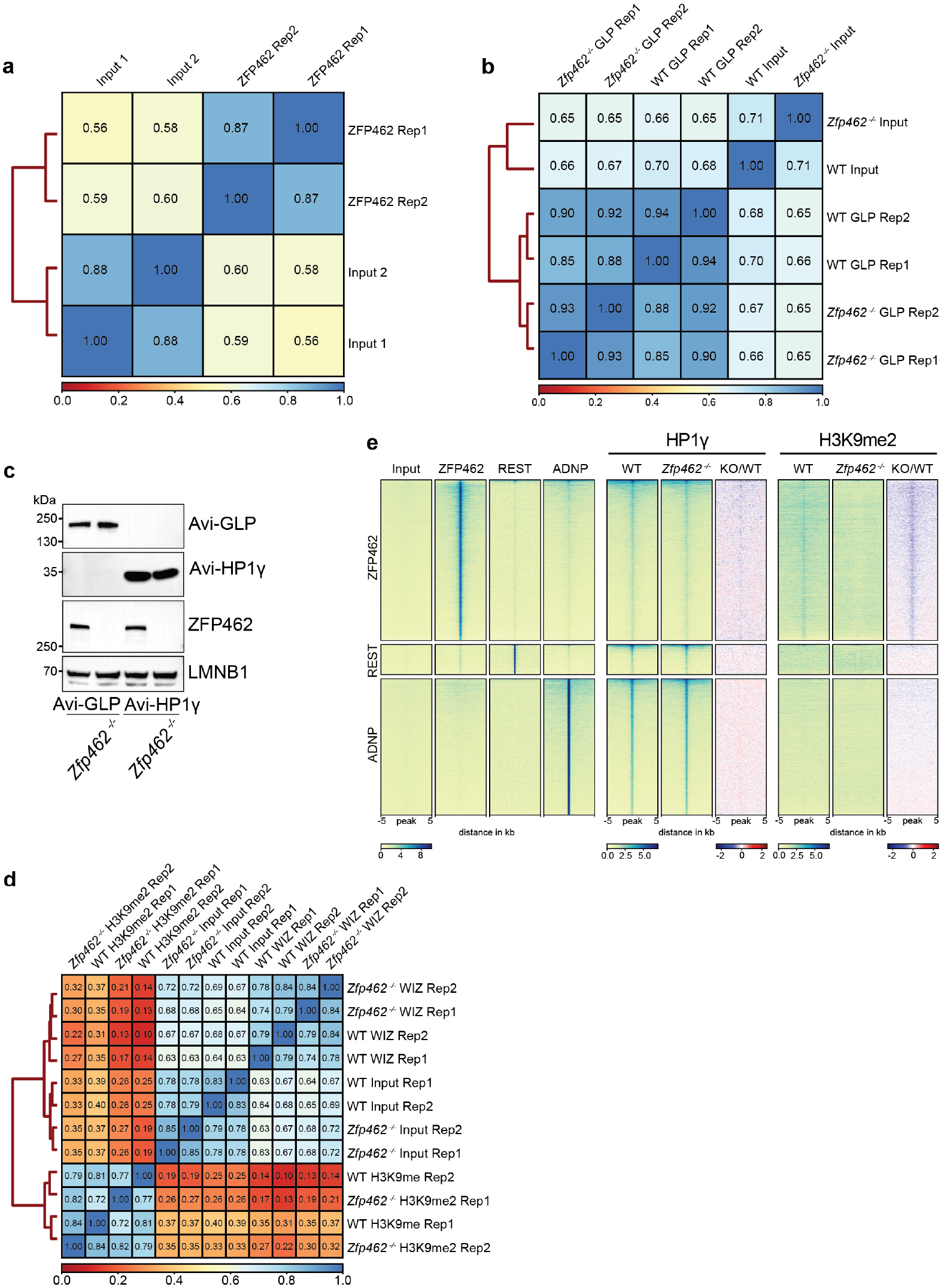
ChIP-seq profiles of ZFP462, REST, ADNP, HP1 and H3K9me2 in WT and *Zfp462* knock-out mESCs. **a)** Heatmap shows Spearman correlation between input controls and two independent ZFP462 ChIP-seq experiments. **b)** Heatmap shows Spearman correlation between input controls and two independent ChIP-seq experiments of GLP and HP1 in WT and *Zfp462 ^-/-^* mESCs. **c)** Western blot shows ZFP462 expression in WT and *Zfp462* Avi-FLAG-tagged GLP and HP1 mESCs. **d)** Heatmap shows Spearman correlation between input controls and two independent ChIP-seq experiments of WIZ and H3K9me2 in WT and *Zfp462 ^-/-^* mESCs. **e)** Heatmaps of ZFP462, REST, ADNP, HP1 and H3K9me2 ChIP-seq signals at ZFP462, REST and ADNP peaks in WT and *Zfp462^-/-^* mESCs. Blue to red scaled heatmap represents ChIP-seq enrichment ratios between *Zfp462^-/-^* versus WT (KO/WT). Each row represents a 10 kb window centred on peak midpoints, sorted by ZFP462, REST and ADNP signal in their respective clusters. Below are scale bars (n = average distribution of two replicates).

**Extended Data Fig. 7.**
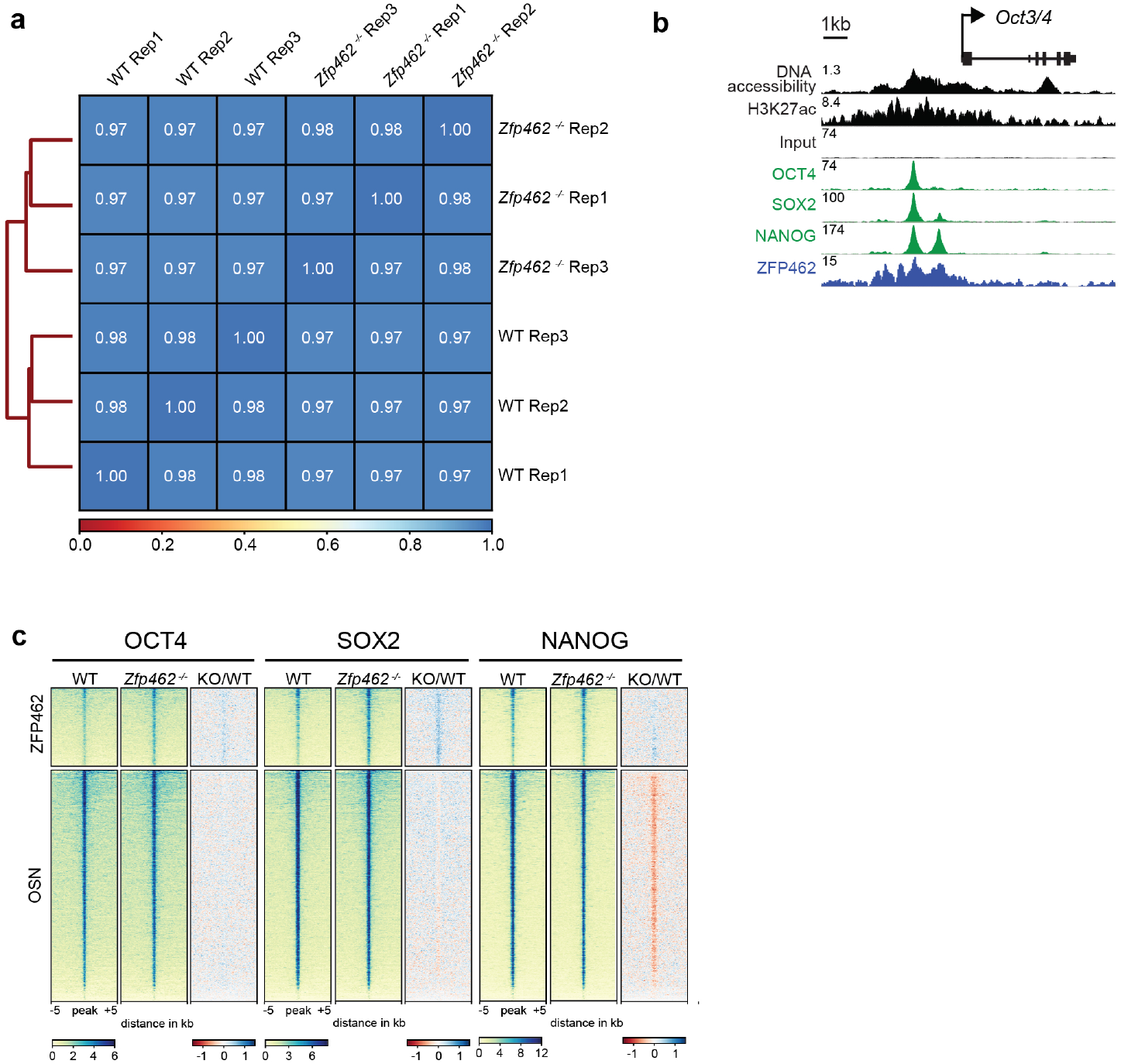
Analysis of chromatin accessibility and genome-wide distribution of pluripotency transcription factors in WT and *Zfp462* knock-out mESCs. **a)** Heatmap shows Spearman correlation between three independent ATAC-seq experiments in WT and *Zfp462* ^-/-^ mESCs. **b)** Genomic screen shot shows DNA accessibility (ATAC-seq) and ChIP-seq signals of H3K27ac, OCT4, SOX2, NANOG and ZFP462 at the *Oct3/4* locus in WT mESCs. ATAC-seq and ChIP-seq profiles are normalized to library size. **c)** Heatmaps of OCT4, SOX2 and NANOG ChIP-seq in WT and *Zfp462 ^-/-^* mESCs at ZFP462-bound peaks with increased DNA accessibility (KO/WT, p-value < 0.05) and OSN peaks. Red to blue scaled heatmap represents ChIP-seq enrichment ratios between *Zfp462^-/-^* versus WT (KO/WT). Each sample row represents a 10 kb window centred on peak midpoints, sorted by wildtype OCT4 ChIP-seq signal intensity (n = average distribution of two replicates).

**Extended Data Fig. 8.**
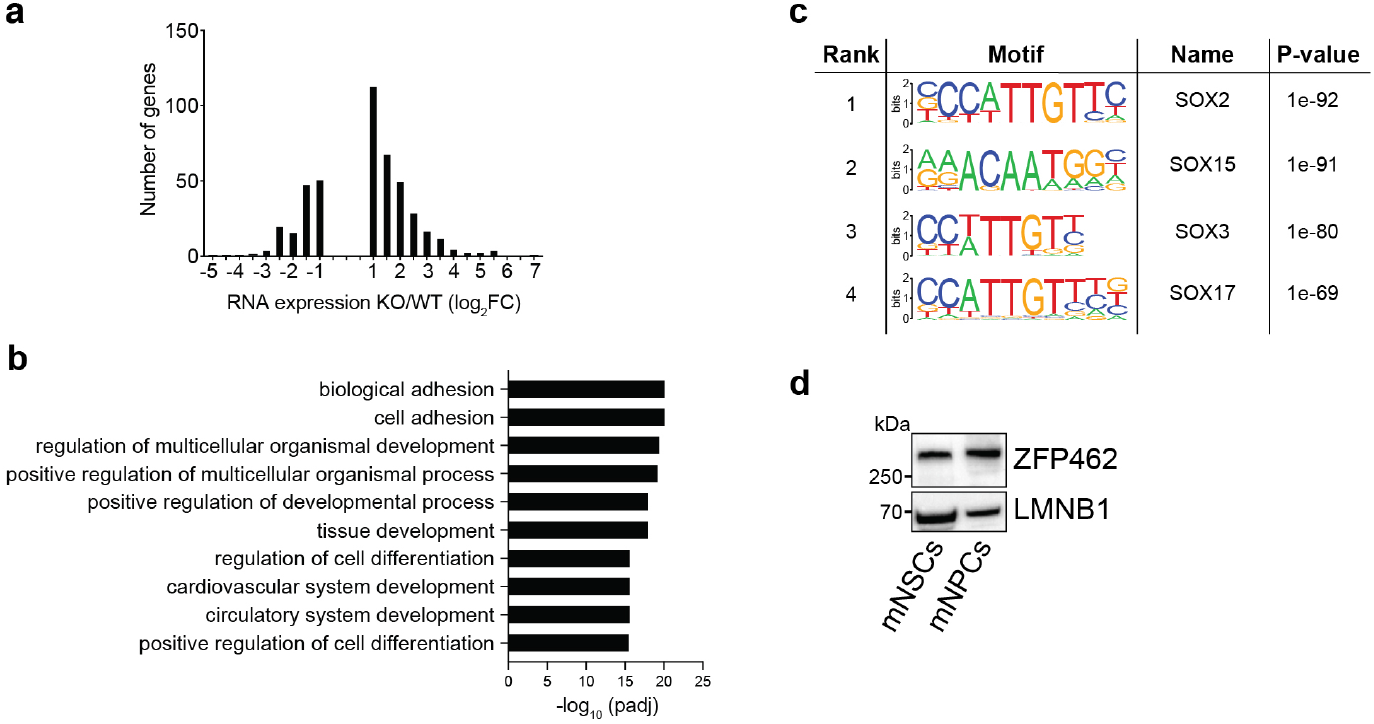
ZFP462-bound enhancers are linked to developmental genes. **a)** Bar plot shows frequency distribution of differential expression for genes located proximal to ZFP462-bound peaks with increased DNA accessibility (KO/WT, p-value < 0.05). **b)** Bar plot shows GO term analysis of biological processes of genes located proximal to ZFP462-bound peaks with increased DNA accessibility (KO/WT, p-value < 0.05). **c)** HOMER analysis of known DNA sequence motifs at ZFP462-bound peaks with increased DNA accessibility (KO/WT, p-value < 0.05). Top ranked DNA sequence motifs and respective significance values are shown in the table. **d)** Western blot shows ZFP462 protein expression in NSCs isolated from mouse brain and NPCs differentiated from WT mESCs.

